# Differentially optimized cell-free buffer enables robust expression from unprotected linear DNA in exonuclease-deficient extracts

**DOI:** 10.1101/2021.09.07.459228

**Authors:** Angelo Cardoso Batista, Antoine Levrier, Paul Soudier, Peter L. Voyvodic, Tatjana Achmedov, Tristan Reif-Trauttmansdorff, Angelique DeVisch, Martin Cohen Gonsaud, Jean-Loup Faulon, Chase L. Beisel, Jerome Bonnet, Manish Kushwaha

## Abstract

The use of linear DNA templates in cell-free systems promises to accelerate the prototyping and engineering of synthetic gene circuits. A key challenge is that linear templates are rapidly degraded by exonucleases present in cell extracts. Current approaches tackle the problem by adding exonuclease inhibitors and DNA-binding proteins to protect the linear DNA, requiring additional time- and resource-intensive steps. Here, we delete the *recBCD* exonuclease gene cluster from the *Escherichia coli* BL21 genome. We show that the resulting cell-free systems, with buffers optimized specifically for linear DNA, enable near-plasmid levels of expression from σ70 promoters in linear DNA templates without employing additional protection strategies. When using linear or plasmid DNA templates at the buffer calibration step, the optimal potassium glutamate concentrations obtained when using linear DNA were consistently lower than those obtained when using plasmid DNA for the same extract. We demonstrate the robustness of the exonuclease deficient extracts across seven different batches and a wide range of experimental conditions across two different laboratories. Finally, we illustrate the use of the Δ*recBCD* extracts for two applications: toehold switch characterization and enzyme screening. Our work provides a simple, efficient, and cost-effective solution for using linear DNA templates in cell-free systems and highlights the importance of specifically tailoring buffer composition for the final experimental setup. Our data also suggest that similar exonuclease deletion strategies can be applied to other species suitable for cell-free synthetic biology.

## INTRODUCTION

Cell-free systems (CFS) have recently revolutionized the field of synthetic biology by providing a fast, simple and efficient platform for rapid genetic-circuit prototyping, biosensor engineering, and decentralized manufacturing ^1^. CFS can be used in basic research for production of recalcitrant or modified proteins ^2^, in educational kits ^3^, for building synthetic cells ^4,5^, and for characterization of biological systems ^6^. They can be deployed for the prototyping of genetic circuits before *in vivo* implementation ^7,8^, such as for metabolic engineering ^9,10^, and also offer an attractive platform for engineering robust, stable and regulatory compliant biosensors with applications in environmental monitoring and medical diagnostics ^11–13^.

Most CFS implementations so far rely on circular plasmids that do not allow utilization of the full potential of the CFS technology. Technically, these setups require DNA cloning and the preparation of large amounts of DNA—processes that can take up to several weeks. Because of the large amount of DNA needed, high-copy number plasmids are preferred for cloning, resulting in high-expression of the gene/s of interest in the cloning strain and subsequent counter-selection in case of metabolic load or cellular toxicity. All these bottlenecks hinder the rapid prototyping capacity afforded by CFS ^14^. The use of linear DNA for cell-free reactions offers the potential to circumvent these issues ^14,15^. When using linear DNA, the cloning step is avoided, and linear expression templates can be simply produced in a few hours by PCR amplification from a plasmid or from synthesized DNA fragments now available at a reasonable cost ^15^. In addition, genes that are toxic in a cellular context can be expressed directly from a linear DNA template. The principal challenge with linear DNA, however, is that it is rapidly degraded by exonucleases naturally present in cellular extracts ^16^. This instability has precluded the widespread use of linear DNA in CFS, and has been limiting the potential of CFS as a rapid prototyping platform.

Recently, several strategies have been developed to enable linear DNA use in CFS. Researchers have started to successfully use the PURE system, a reconstituted expression system made of purified recombinant proteins and devoid of exonuclease activity ^17^. Yet, the PURE system is expensive and does not support expression from native promoters. The ROSALIND system takes a different approach to the engineering of cell-free biosensors but lacks the translation step ^18^. To achieve linear DNA expression specifically in cell-extract based CFS, many protection strategies have been developed. These include using the λ-phage GamS protein ^19^ or DNA containing Chi sites ^20^ to inhibit the RecBCD exonuclease complex. Protecting the linear DNA ends by DNA binding proteins has also been successful ^21–23^, as has the chemical modification of linear DNA ends ^14^. All these methods still require supplementation of the extract with additional time- and cost-intensive components or modification of the template to achieve expression from linear DNA. Furthermore, reagents such as GamS can be costly, complicating their use at large scale. Adding supplementary reagents also limits the working volume of the reactions, and can be the source of unwanted experimental variation that reduces the reproducibility of cell-free reactions.

Here we take an alternative approach to stabilize linear DNA in CFS. Instead of protecting the ends of the linear DNA template or inhibiting the exonucleases, we knockout the exonuclease genes from the genome before preparing the cell extract. The exonuclease V complex RecBCD has long been known to degrade linear DNA templates in cell-free extracts ^16^. Previous work also showed that *E. coli* extracts from a RecB mutant strain and a Δ*recBCD* stain resulted in stabilization of and expression from linear DNA templates ^24,25^. Those works were done in strains different to the now commonly used CFS chassis, (*E. coli* BL21 and its derivatives), and the more recent papers have achieved high expression by using the T7 promoter ^25,26^. Nevertheless, these papers demonstrated that disruption of the *recBCD* operon from the chassis strain can offer an elegant solution to the problem of linear DNA templates’ degradation. Yet, the exonuclease deficient A19 strain has not been extensively adopted by the CFS community for expanding the use of linear DNA templates. A19 extract is reported to have lower productivity than extracts from BL21 (by ∼30%) and BL21 Rosetta (by ∼43%) strains for cell-free protein production ^19,27^.

In this work, we deleted the *recB* gene or the full *recBCD* operon from *E. coli* BL21 Rosetta2, optimized their cell extracts specifically for linear DNA expression from native *E. coli* promoters, and deployed them for characterization of toehold switches and enzyme variants. Unlike what has been previously reported ^25,28^, neither of these deletions had a major effect on cell growth or viability. The resulting extracts support expression from linear DNA templates without the addition of any extra component or DNA template modifications. We show that buffer optimization specifically for linear DNA expression improves protein production yield from native σ70 promoters without the use of GamS protein or Chi DNA, and that purification of the PCR products can also be avoided. These results are valid across two different laboratories for different experimental conditions (lysis method, DNA concentration, temperature, GFP gene, flanking DNA length, DNA purification, plate type, and plate reader). The buffer-specific differences in gene expression are not due to differences in DNA stability, but probably due to a transcription-related mechanism. Finally, we demonstrate that Δ*recBCD* extracts support several applications such as rapid screening of toehold switches and activity assessment of enzyme variants. Our work provides a simple, efficient, and cost-effective solution for using linear DNA templates in *E. coli* cell-free systems. We anticipate that Δ*recBCD* cell extracts will facilitate protein production and rapid prototyping of genetic circuits in CFS for metabolic engineering, biosensing, and manufacturing.

## RESULTS

### Exonuclease deficient *E. coli* BL21 extracts exhibit modest expression from linear DNA

Due to its superior protein production ability, the *E. coli* BL21 strain and its derivatives have been a workhorse of protein production applications for decades. More recently, *E. coli* BL21 strain ^21,22,29^, and its Rosetta2 derivative ^6,30–33^, have also been widely adopted for use in the production of *E. coli* extracts for cell-free systems. In this work, both these strains were used for cell-free extract preparation (**Sup Tables 1** and **2**), which also allows us comparison with other linear DNA papers ^20–22,33^. Inspired by previous work where knocking out or mutation of *recBCD* genes in *E. coli* prior to extract preparation increased protection of linear DNA ^24,25^, we used a two-step λ-Red recombination protocol (**Methods**, ^34^) to generate Δ*recB* and Δ*recBCD* knockouts from the *E. coli* BL21 Rosetta2 (**Fig 1A**). The KO strains were confirmed by PCR (**Sup Fig 1**) and sequencing (**Fig 1B**). Δ*recB* mutation has previously been reported to substantially reduce fitness of *E. coli* ^25,28^. However, we found less than 25% reduction in specific growth rates in the Δ*recB* and Δ*recBCD* strains when compared to their respective parental strains (**Sup Fig 2**). This comparison was performed for both the BL21 strains lacking the pRARE2 plasmid and the BL21 Rosetta2 strains containing the pRARE2 plasmid. pRARE2 overexpresses low-frequency tRNAs in the Rosetta2 strain to allow testing of protein coding genes not specifically optimized for *E. coli*^35^.

**Figure 1:**
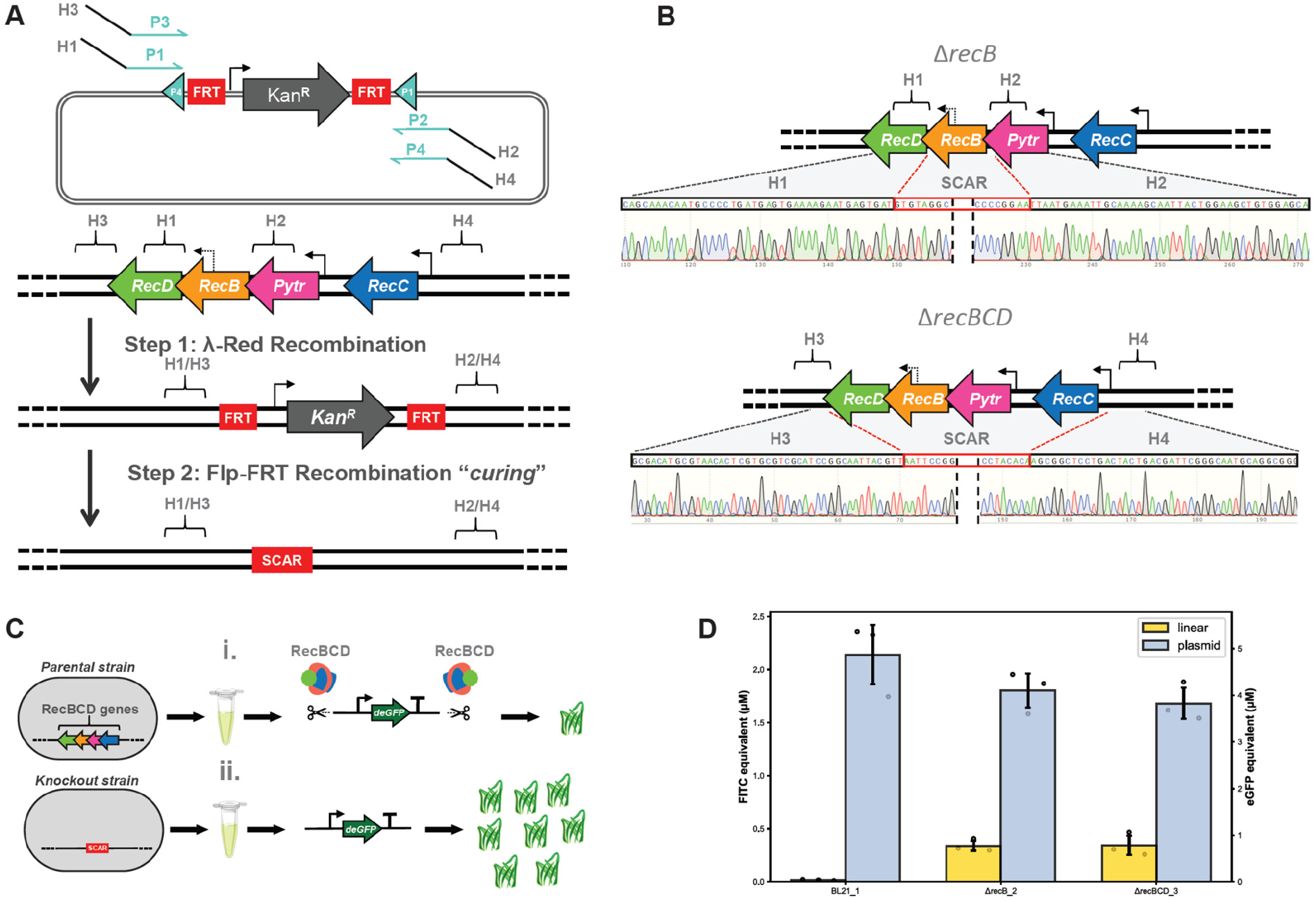
Cell-free extracts from Δ*recB* and Δ*recBCD E. coli* BL21 support modest expression from linear DNA. **(A)** *E. coli* genome was edited using λ-Red recombination. The pKD13 plasmid was used as a template to amplify Kanamycin resistance cassette (*Kan*^*R)*^ using primers containing 5′-tails H1 & H2 or H3 & H4, homologous to the *recB* or the *recBCD* locus. The loci were deleted by replacement with *Kan*^*R*^ cassette and later cured using the Flp-FRT system. **(B)** Gene deletions were verified by Sanger sequencing after excision of the *Kan*^*R*^ cassette from previous homologous recombination. **(C)** Linear DNA is generally degraded in *E. coli* extracts by the RecBCD holoenzyme, preventing high gene expression. Deletion of *recBCD* allows linear DNA stability and, consequently, high gene expression. **(D)** Linear DNA (5 nM) expression is near zero in BL21 Rosetta2 (BL21_1) extract in comparison to equimolar plasmid DNA. However, upon disruption of either *recB* (ΔrecB_2 extract) or *recBCD* (ΔrecBCD_3 extract) modest linear DNA expression is observed. Buffer compositions for the three extracts were previously optimized using 1 nM plasmid DNA. Data shown are the mean±SD for 3 replicates done on the same day. Additional experimental conditions are listed in Sup Table 6.

The *E. coli* BL21 Rosetta2 WT, Δ*recB*, and Δ*recBCD* strains were used for cell-free extract preparation and buffer calibration using plasmid DNA (**Methods**), and expression from equimolar (5 nM) amounts of plasmid and linear *deGFP* DNA compared across extracts (**Fig 1C, 1D**, and **Sup Fig 3**). Linear DNA is unprotected in WT extracts, producing <1% fluorescence compared to plasmid DNA in the same extract. However, Δ*recB* and Δ*recBCD* extracts result in modestly higher GFP fluorescence from the linear DNA at 19% and 21% of plasmid DNA expression levels, respectively. These expression levels are lower than those reported from *E. coli* A19 ^25^, where exonuclease V deletion restored expression from linear DNA to 64% of plasmid levels. A key difference is that our *deGFP* transcription is driven by the native σ70 promoter while many of the previous linear DNA strategies have used a T7 promoter to achieve higher expression in CFS ^25,26^. Other works that used native σ70 promoters for linear DNA expression have also reported lower expression from linear DNA ^33^.

### Differential optimization of exonuclease-deficient extracts substantially improves cell-free expression from linear DNA

Consistent with findings from other labs ^8,22,36,37^, we have observed considerable variation in total GFP expression levels from one batch of CFS to another. In our previous work, we improved expression from *E. coli* cell-free extracts by combinatorial buffer optimization for 11 components, where buffer-specific optimization was able to reduce batch-to-batch variation in extract productivity ^36^. Given the critical role buffer composition plays in CFS, many studies have optimized it specifically for each plasmid used ^30,38^. Therefore, we wondered if the optimal buffer composition for linear and plasmid DNA expression in CFS should also be differentially calibrated. Starting with a standard laboratory protocol for cell-free extract preparation from BL21 (-pRARE2) strains using French press lysis (**Methods**), the buffer composition for those extracts was calibrated separately for linear and plasmid DNA. The buffers were sequentially optimized first for Mg-glutamate and then for K-glutamate levels (**Fig 2A**). We find that the optimal Mg-glutamate levels for both linear and plasmid DNA are similar across two batches of Δ*recBCD* extracts at 7-8 mM (**Fig 2B**). However, the optimal K-glutamate levels for *sfGFP* expression from linear and plasmid DNA are substantially different in the same extract. The optimal K-glutamate levels for plasmid DNA are 160 mM and 220 mM for the two extract batches (ΔrecBCD_5 and ΔrecBCD_6), whereas the optimal K-glutamate levels for linear DNA are 80 mM and 140 mM for the same batches, respectively (**Fig 2C**). The DNA concentration needed for optimal GFP expression is ∼10 nM for both the WT and Δ*recBCD* extracts (**Sup Fig 4**).

**Figure 2.**
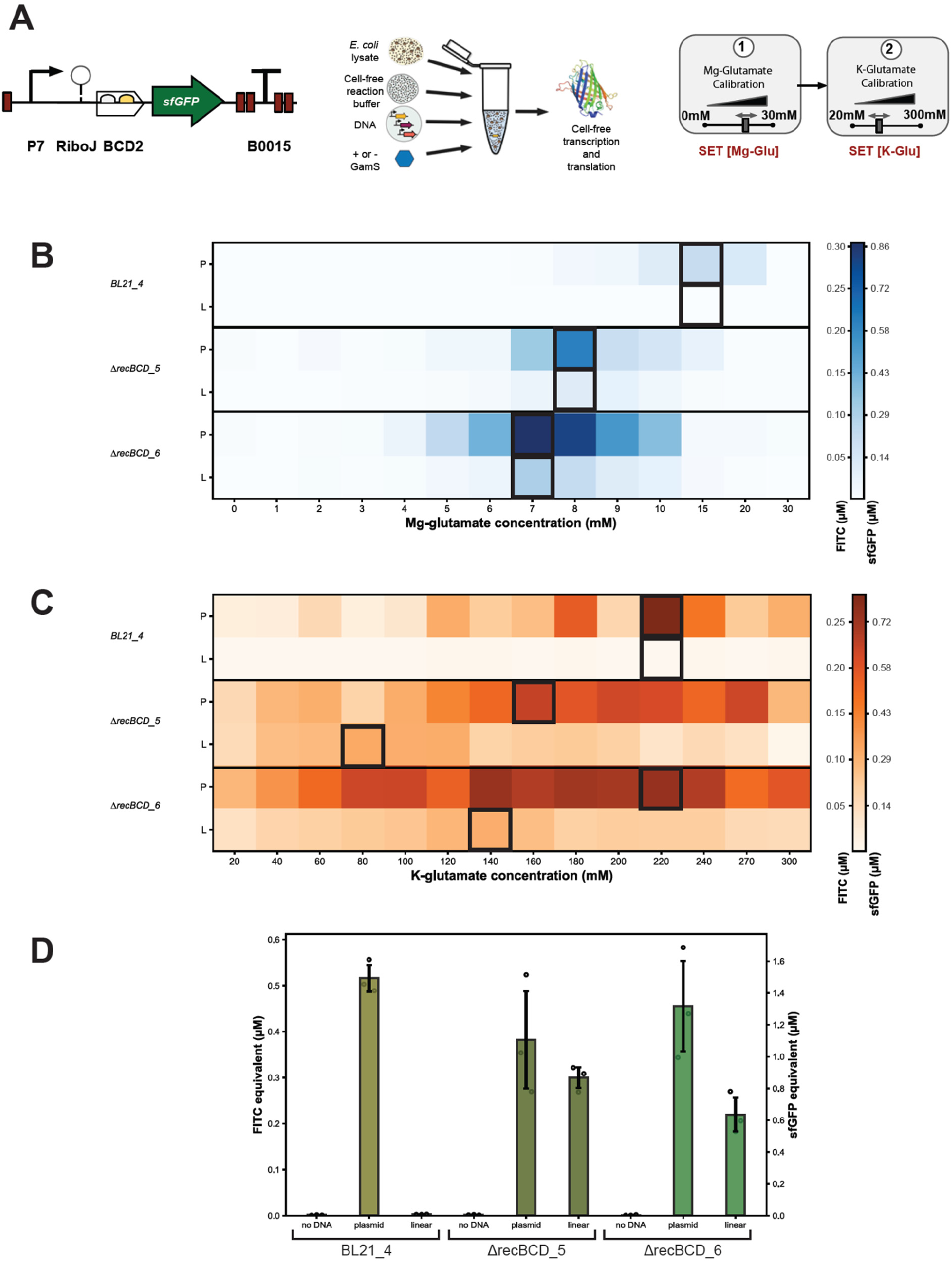
Specific cell-free calibration improves gene expression from linear DNA template. **(A)** Diagram of sfGFP under the control of the strong P7 constitutive promoter, with the self-cleaving ribozyme RiboJ and the bicistronic RBS BCD2. The DNA amplified by PCR was used as a template to calibrate the reaction buffer of the cell-free system. **(B)&(C)** Heatmaps of the sequential Mg-glutamate calibration (first step, with 100 mM K-glutamate) and K-glutamate calibration (second step, with the optimal Mg-glutamate concentrations) of three different batches of extracts with a plasmid DNA template (P) or a linear fragment (L) from the same expression unit. BL21_4 stands for the WT BL21 extract, and ΔrecBCD_5 and ΔrecBCD_6 for the two batches of extracts from the Δ*recBCD* BL21 strain. sfGFP expression was measured after 8 hours at 37 °C in 20 µL reactions. DNA concentration was fixed at 4 nM for both plasmid and linear templates. Boxes highlight the optimal concentration of Mg-glu or K-glu for each strain and expression template. Data shown in the heat-maps are from a single experiment. **(D)** 8-hour time point fluorescence from experiments on three different days for the three calibrated extracts. Linear and plasmid DNA were added to a final 10 nM concentration in a 20µL reaction (see Sup Fig 7). Data shown are mean±SD for 3 replicates done on different days. Additional experimental conditions are listed in Sup Table 6.

Previous studies ^37^, including from our own lab ^36^, have shown considerable variability in cell-free expression across personnel and laboratories. To ascertain whether our unexpected results hold across a variety of conditions, the differential buffer calibration was repeated in a different lab for three additional batches of exonuclease-deficient extracts prepared using sonication lysis of BL21 Rosetta2 strains (+pRARE2) and calibrated using a different gene (*eGFP*) encoded on linear and plasmid DNA (**Methods, Sup Fig 5**). We find that the optimal Mg-glutamate levels for both linear and plasmid DNA are similar across Δ*recB* and Δ*recBCD* extracts at 7-8 mM (**Sup Fig 5B**). However, as before, the optimal K-glutamate levels for plasmid DNA for the three extracts are 140-160 mM, whereas the optimal K-glutamate levels for linear DNA for the same extracts are 20-40 mM (**Sup Fig 6C**). By analysing seven different extracts (two WT, and five Δ*recB*/Δ*recBCD*) prepared using different parental strains (-pRARE2 and +pRARE2) and lysis methods (French press and sonication), calibrated using two different genes (*sfGFP* and *eGFP*), and incubated at two different temperatures (37 °C and 30 °C), we find that buffer composition of cell-free extracts must be differently calibrated for optimal expression from linear and plasmid DNA used in exonuclease V deleted extracts.

Purification of PCR products before running the linear DNA cell-free reactions adds cost and time limitations to rapid prototyping. Additionally, a PCR purification step can result in the loss of ∼50% DNA. So, we investigated whether purification of PCR products can be avoided. Comparable expression of GFP was found in cell-free reactions with 10 nM of purified and unpurified PCR-amplified linear DNA (**Sup Fig 6**), where the purified DNA was quantified by Nanodrop and the unpurified DNA by Qubit (**Methods**). These results allow us to avoid the time-consuming purification step entirely.

Next, we compared the expression levels from equimolar amounts of linear and plasmid DNA (10 nM) in the WT and the two batches of Δ*recBCD* extracts (**Fig 2D**). sfGFP expression from linear DNA reached 48-78% of that from plasmid DNA in the two batches of the Δ*recBCD* extracts prepared using French press lysis. These results also hold true for eGFP expression in Δ*recBCD* and Δ*recB* extracts prepared using sonication lysis (**Sup Fig 5D**). Expression from linear DNA reaches 101-138% of that from plasmid DNA in these Δ*recB* and Δ*recBCD* extracts. Together, these results show that expression from linear DNA, driven by a native *E. coli* σ70 promoter, can be increased to near-plasmid levels when the cell-free buffer is specifically calibrated for linear DNA.

### Increased linear DNA expression in low K-glutamate buffer is not due to differences in DNA degradation

Across the seven extracts tested above, the optimal Mg-glutamate concentrations were found to be between 7-9 mM. The optimal K-glutamate concentrations lie in a broader range of 140 to 240 mM, when using plasmid DNA for calibration. These values are broadly consistent with the optimal values obtained in other studies that also used plasmids for buffer calibration ^30,36,39^. Surprisingly, the optimal K-glutamate concentrations obtained above when using linear DNA for buffer calibration were between 20-140 mM, and always lower than the optimal K-glutamate concentrations obtained when calibrating the same extract using plasmid DNA (**Fig 2, Sup Fig 5**). We wondered if K-glutamate concentrations affect the activity of other *E. coli* nucleases that may be present in the cell-free extract, especially exonuclease IX whose DNA binding function is K^+^ dependent ^40^. To investigate whether the increase in linear DNA expression in the low K-glutamate buffer was due to a decrease in DNA degradation, we ran linear DNA cell-free reactions in *E. coli* BL21 Rosetta2 WT, Δ*recB*, and Δ*recBCD* extracts with both low and high K-glutamate buffers (**Fig 3A**), isolated DNA from them at three time-points (at 0, 1, and 8 hours), and quantified the DNA by qPCR (**Methods**). Unsurprisingly, >99% of the linear DNA is lost within 1 hour in the WT extracts irrespective of the buffer used (**Fig 3B**). In contrast, the loss of DNA in the Δ*recB* and Δ*recBCD* extracts by 8 hours is 29-54%. This confirms that *recBCD* deletion decreases linear DNA degradation. However, there is no clear difference in the degradation observed between the two buffer compositions, suggesting that a mechanism other than DNA degradation is responsible for increased expression in the low K-glutamate buffer.

**Figure 3.**
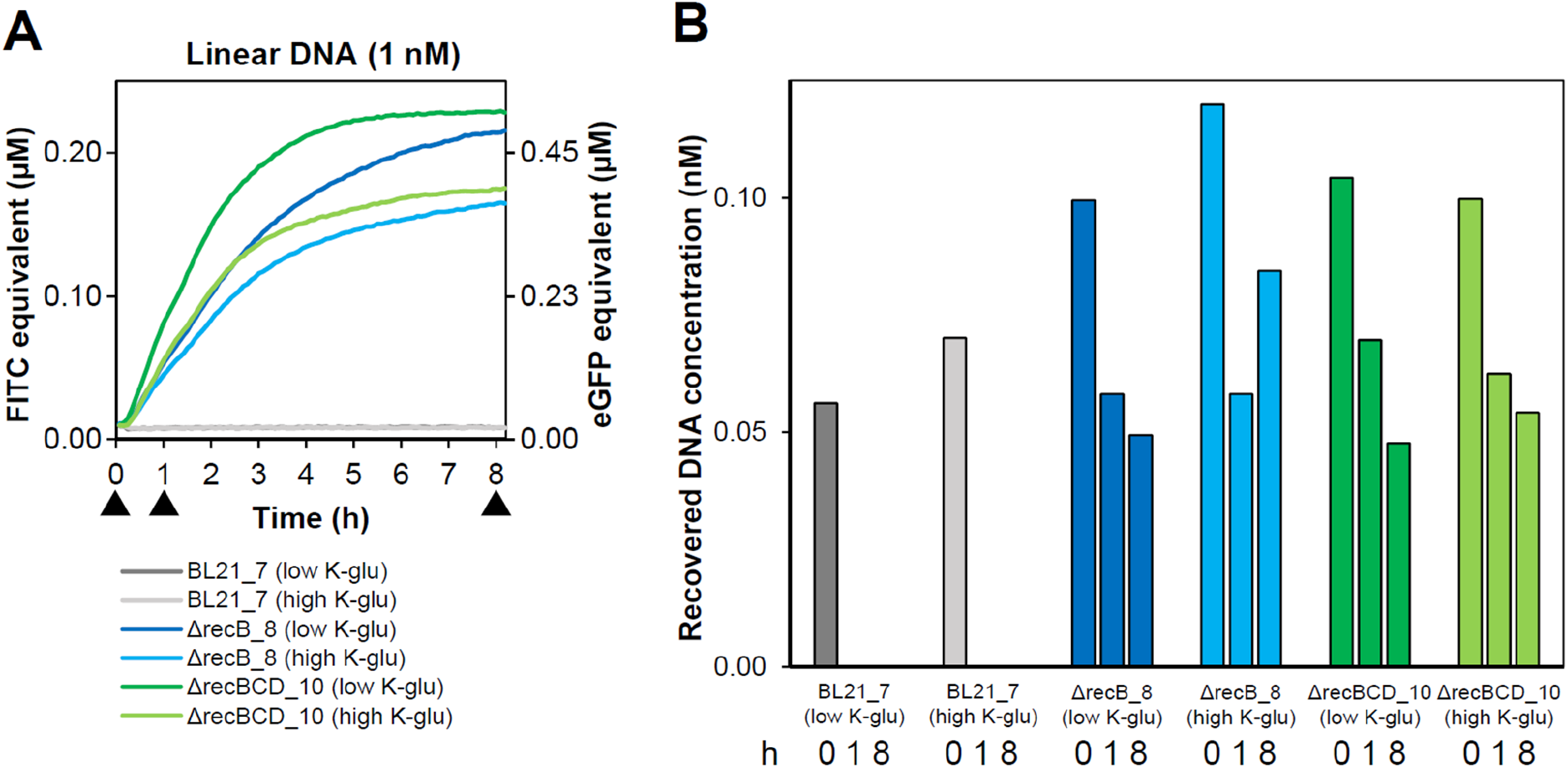
Investigating buffer-specific differences in gene expression from linear DNA in CFS. **(A)** GFP expression was monitored over ∼8 hours from linear DNA in cell-free extracts from *E. coli* BL21 Rosetta2 WT, Δ*recB* and Δ*recBCD* strains, in two different buffers (low K-glutamate and high K-glutamate). The runs were interrupted at three time-points (0, 1, and 8 hours) to extract residual linear DNA from the CFS. Data shown are kinetic plots from a single cell-free experiment. **(B)** Measurement of DNA concentration from each reaction at the three time-points by quantitative PCR. Data shown are mean DNA concentrations from 3 qPCR technical replicates performed on DNA recovered from a single cell-free experiment (Methods). Additional experimental conditions are listed in Sup Table 6.

It has been previously shown that high concentrations of K-glutamate reduce the binding of *E. coli* RNAP to DNA ^41^. Increasing K-glutamate concentrations from 50 mM to 300 mM was found to abolish RNAP binding to both plasmid and linear DNA, in turn reducing cellular transcription ^42^. It is possible then that a low K-glutamate buffer supports higher GFP expression due to increased transcription from a native σ70 promoter, a buffer-specific difference that is uncovered only when using the relatively unstable linear DNA templates.

### Various linear DNA protection methods do not improve protein production in exonuclease-deficient extracts

Encouraged by our results above and previous work that has shown that strategies for linear DNA protection can have an additive effect ^43^, we further tested whether combining exonuclease-deficient cell extracts with previously reported methods of linear DNA protection increases GFP expression even further (**Fig 4**). As our data show that the Δ*recBCD* extracts exhibit slightly higher GFP expression from linear DNA than the Δ*recB* extracts (**Fig 3A, Sup Fig 5D**), we decided to henceforth work only with the Δ*recBCD* extracts.

**Figure 4.**
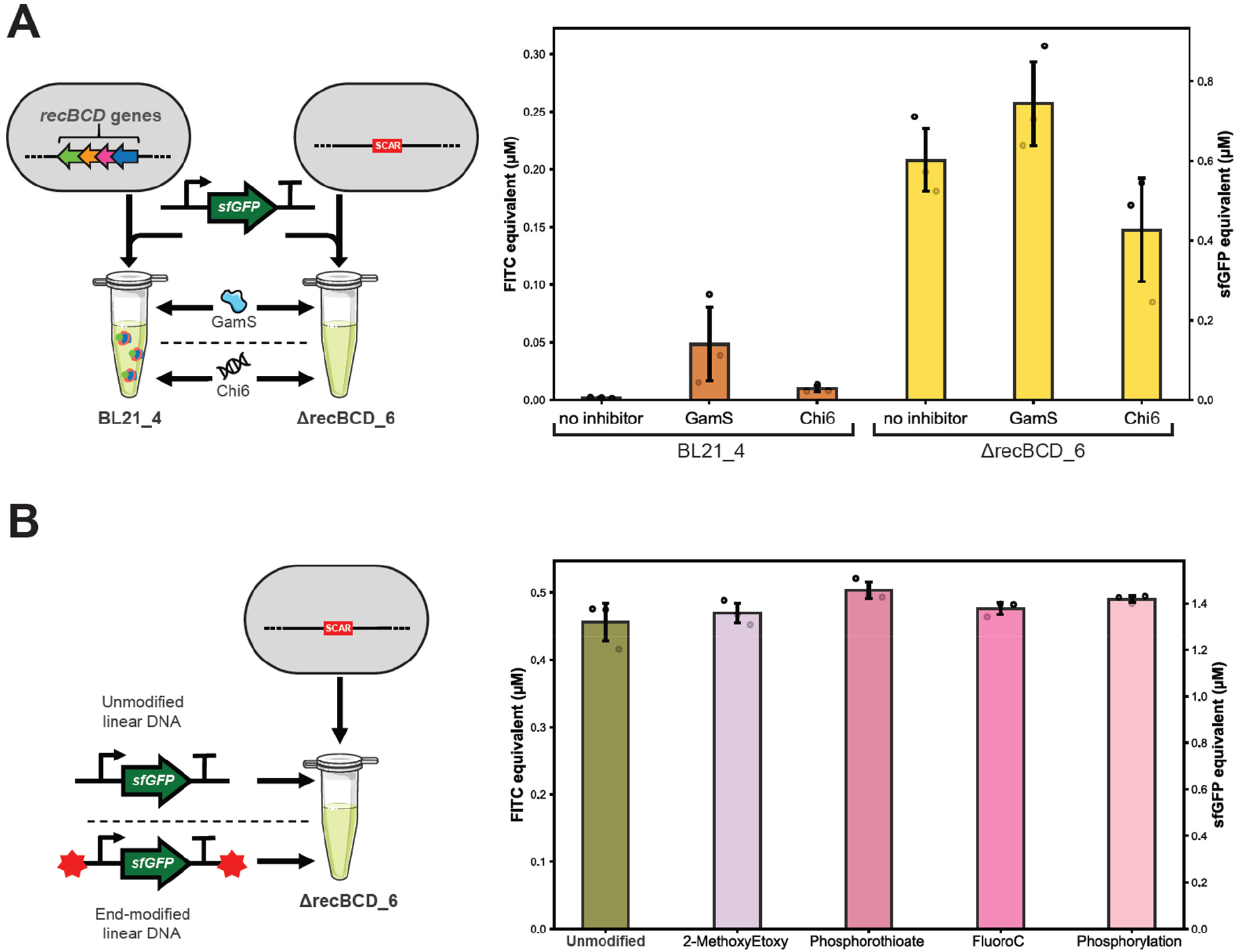
Effect of various linear DNA protection methods on protein production in *ΔrecBCD* extracts. **(A)** Comparison of sfGFP expression with GamS and Chi6 nuclease inhibitors. GamS protein and Chi6 oligonucleotides were added at a final concentration of 2 µM (see Methods and Sup Fig 7). Data shown are mean±SD for 3 replicates done on different days. **(B)** Assessment of the effect of various chemical modifications on the 3′ and 5′ ends of the linear DNA fragment on gene expression in ΔrecBCD_6 extract. The reporter cassette was PCR amplified using various modified oligonucleotides and used as a template in cell-free reactions (Methods). Data shown are mean±SD for 3 replicates done on the same day. Additional experimental conditions are listed in Sup Table 6.

GamS not only inactivates the RecBCD exonuclease but also other nucleases like SbcC by blocking their catalytic sites ^44,45^. So, we investigated whether a combination of *recBCD* deletion and GamS-mediated sequestration of any other residual nucleases may further improve linear DNA expression in our extracts. To test if supplementation with GamS increases linear DNA protection, purified GamS protein was first produced using an in-house protocol (**Methods**), and then the concentration of GamS needed for optimal linear DNA expression was identified in our extract to be 1.5 μM (**Sup Fig 7A**). Next, we compared GFP expression from linear DNA in both WT and Δ*recBCD* extracts (**Fig 4A**) and found that although GamS supplementation improves linear DNA expression in the WT extract it does not substantially improve linear DNA expression in the Δ*recBCD* extract. While GamS supplementation allows us to precisely calibrate the optimal GamS concentrations needed for linear DNA protection, it is expensive to buy or purify in-house. The supplementation also leads to the loss of 1 µL out of the total 5 µL working volume available in a 20 µL cell-free reaction (the remaining 15 µL is occupied by the cell-free lysate and buffer). Therefore, we further investigated if the results obtained using purified GamS can be reproduced using GamS-doped extracts. GamS expression in *E. coli* cells was first induced, followed by extract preparation and differential calibration of the extracts for linear and plasmid DNA (**Methods, Sup Fig 8B** & **8C**). We compared linear DNA expression in GamS-doped WT, Δ*recB*, and Δ*recBCD* extracts (**Sup Fig 8**). The results confirmed our previous findings that GamS supplementation increases linear DNA expression in WT extracts but does not substantially increase expression in exonuclease V deleted extracts (Δ*recB* and Δ*recBCD*) (**Sup Fig 8D**). GamS induction prior to extract preparation seems to reduce linear DNA expression in the Δ*recBCD* extract by 36%. Surprisingly, GamS-induced extracts of all three genetic backgrounds (WT, Δ*recB*, and Δ*recBCD*) also exhibit reduced plasmid DNA expression. This could be due to altered physiology during growth on ampicillin selection and/or induction of GamS by arabinose prior to extract preparation.

Chi DNA (χ-DNA) has been shown to increase linear DNA expression by sequestering the RecBCD complex ^20^. We tested whether adding χ-DNA to our Δ*recBCD* extracts will increase linear DNA expression further. DNA with 6 χ-repeats per double-stranded molecule (Chi6) was assembled and the concentration of χ-DNA needed for optimal linear DNA expression in our extract was identified to be 0.75 μM (**Sup Fig 7B**). Next, we compared GFP expression from linear DNA in both WT and Δ*recBCD* extracts (**Fig 4A**) and found that similar to GamS, χ-DNA only modestly increases linear DNA expression in WT extracts but it does not increase linear DNA expression in Δ*recBCD* extracts. Taken together, our findings from the GamS and χ-DNA supplementation experiments confirm that the RecBCD complex is the primary driver of linear DNA degradation in *E. coli* extracts, and, together with buffer calibration, its deletion provides sufficient protection to linear DNA.

Terminal protection of the linear DNA can also be achieved by chemically modifying the DNA ends. DNA methylation (Zhu et al, 2020) and terminal phosphorothioates (Sun et al, 2013) were shown to improve expression by 32% and 36% ^14^, respectively. We hypothesized that other chemical modifications known for protecting against exonucleases degradation could further increase the yield in our RecBCD-deleted system. The ability of different modifications on DNA ends to confer additional protection for linear DNA expression in our Δ*recBCD* extracts was tested. We evaluated four modifications: 2-methoxyethoxy, phosphorothioate, fluoroC, and phosphorylation (**Sup Table 3**). No significant difference in linear DNA expression was found in the Δ*recBCD* extracts from any of the end-modified linear DNAs (**Fig 4B**). This suggests that these modifications do not substantially affect linear DNA protection in the Δ*recBCD* extracts. In the exonuclease-containing WT extract, only the phosphorothioate modification increased expression from linear DNA templates to 2-fold (**Sup Fig 9A**). However, the overall expression remained much lower than that from plasmid DNA, or that in exonuclease-deficient extracts (**Fig 4B, Sup Fig 9B**). These data suggest that exonuclease depletion from the cell-free extracts confers higher protection to linear DNA (in the appropriate buffer) than chemical modification of DNA ends.

### Rapid testing of toehold switches

RNA fulfils several regulatory, enzymatic, and structural roles in cells, many of which can be used to reprogram cellular functions ^46^. Specifically, RNAs have been engineered as biosensing devices for many small molecules and other RNAs ^47^. A subclass of RNA-sensing riboregulators are the toehold switches that combine modular architecture with secondary structural constraints to generate readily programmable sensors for an arbitrary RNA sequence^48^. Toehold switches, or their SNIPR variants, have already been deployed in CFS for the sensing of cancer mutations, Ebola virus, Zika virus, and norovirus ^13,49–51^. While the required secondary structural constraints of toehold switches are broadly understood, *de novo* design of toehold switches remains a challenge due to the many unknowns that define their sequence-structure-function relationship, recently prompting the use of machine learning methods for improved design ^52,53^. Despite these efforts, several toehold switch designs typically need to be experimentally validated to identify the best candidate ^53^.

Having established above a method for robust expression from unprotected linear DNA, we next used the method to demonstrate rapid screening of toehold switch variants for the previously published gen2_num01 trigger RNA from Green et al. ^48^, here expressed from the plasmid sTR060. The toehold switch consists of an unpaired toehold region at the 5′-end, whose initial interaction with the trigger RNA unfolds the switch, followed by a stem-loop structure that in the OFF state prevents translation initiation at the start codon (**Fig 5A**). In this architecture, the ribosome binding site (RBS) is located in the unpaired loop region and determines the rate of translation in the ON state of the switch. The sTR056 toehold switch was derived from the original gen2_num01 switch ^48^ and its variants generated with the original RBS (CAGAGGAGA) as well as 11 other sequences (**Fig 5B, Sup Fig 10**). Each switch was PCR amplified as linear DNA and directly added to the Δ*recBCD* CFS (**Methods**), with or without the pre-transcribed trigger RNA.

**Figure 5.**
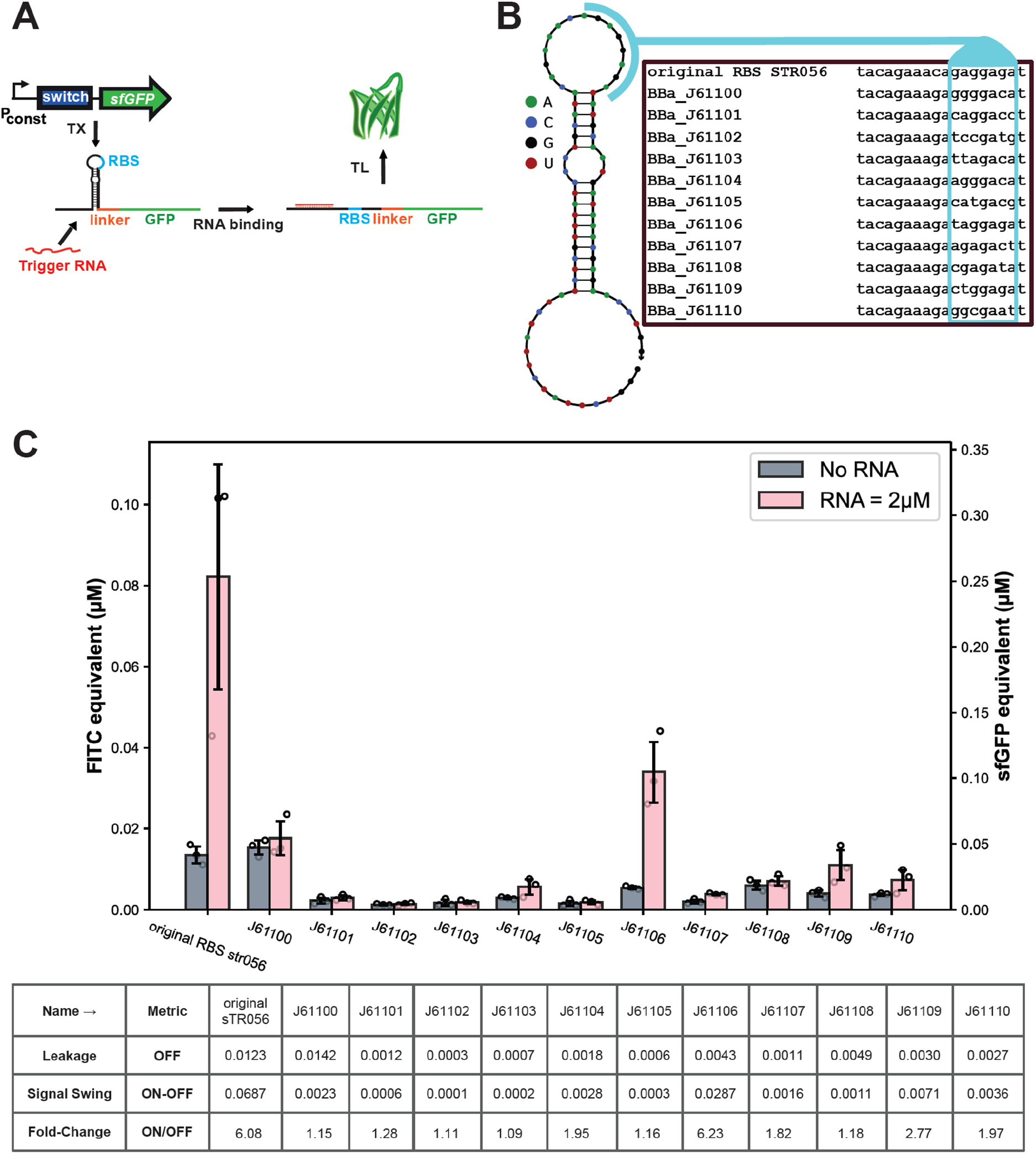
Rapid assessment of toehold-switch function using linear DNA in Δ*recBCD* extracts. **(A)** Principle of RNA-sensing using toehold switches. The reporter mRNA is designed with a hairpin loop preceding the coding sequence and blocking translation. This hairpin is designed to be unfolded by pairing with a specific trigger RNA that restores translational activity. **(B)** Toehold-switch library with variable RBS design. In order to modulate the activity of a model toehold switch, variants were designed by changing the RBS using the sequences from the iGEM registry/ Anderson library (see Methods). The blue box depicts the region of sequence variation. (**C**) Linear DNA mediated cell-free characterization of toehold switches variants. 10 nM of linear DNA encoding sensor candidates were expressed in Δ*recBCD* cell-free extract in presence or absence of 2 µM of trigger RNA. Variants with new characteristics of leakage, signal swing, and fold-change were identified. Data shown are mean±SD for 3 replicates done on different days. Additional experimental conditions are listed in Sup Table 6.

The results showed a variety of switch behaviours in the library (**Fig 5C**), with leakiness between 0.3 and 14.2 nM FITC equivalents, signal swing between 0.1 and 68.7 nM FITC equivalents, and fold-change between 1.09 and 6.23. The J61106 RBS variant shows a fold-change comparable to the original RBS, but has a lower leakiness in the OFF state. We demonstrate for the first time the rapid screening of toehold switch designs using linear DNA in extract-based cell-free systems while using native *E. coli* promoters for switch expression. Our inexpensive approach requires neither the PURE cell-free system, GamS or Chi DNA supplementation, chemical modification of DNA ends, nor T7 promoters or polymerase for switch expression.

### Screening of enzymes using linear DNA

Riboswitches and transcription factors (TFs) can be used in CFS for biosensing of small molecules in clinical or environmental samples ^11,54,55^. Usually, this is only possible if the molecule of interest is a ligand for a known RNA aptamer or TF. However, we have shown that sensing-enabling metabolic pathways (SEMPs) can also be used in CFS to convert undetectable molecules into detectable ones using transducer enzymes ^12,56^. Conversely, biosensing TFs can be used for monitoring upstream enzymatic activity for biotechnological applications ^29^. To assess the capabilities of our newly developed linear DNA system, it was applied to a classical problem in biotechnology: the rapid screening of recombinant enzyme activity. In fact, identifying enzyme candidates with good cell-free expression and activity levels is a key bottleneck for the development of new biosensors and the construction of new metabolic pathways for bioproduction of compounds of interest.

Hippurate hydrolase (HipO) is an enzyme produced by several bacteria that catalyses the following reaction:

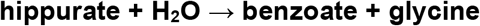

HipO has been used as a metabolic transducer allowing the detection of hippurate, which upon conversion to benzoate binds and activates the transcription factor BenR that, in turn, activates a reporter gene (**Fig 6A**) ^12^. This cell-free optimized benzoate biosensor can be used to monitor benzoate production in the system, thereby enabling the screening of different HipO enzyme variants. We identified new HipO homologues from four bacterial species (*Salmonella enterica, Helicobacter felis, Helicobacter ailurogastricus*, and *Campylobacter coli*) (**Methods**). All of these enzyme variants share at least 90% amino acid sequence identity (**Fig 6B, Sup Fig 11**). Next, we used our linear DNA CFS to compare their activities against the previously tested *Campylobacter jejuni* HipO. Codon-optimized genes encoding the enzyme variants were synthesized as full-length expression cassettes and added to the cell-free reaction in the form of PCR-amplified linear DNA fragments, whereas the TF and reporter genes were expressed from plasmids (**Fig 6C**). Cell-free expression of these enzyme candidates showed that out of the four new variants, two exhibited hippurate to benzoate conversion levels at least similar or better than the reference sequence from *C. jejuni*, while one showed a weak conversion level, and the last one demonstrated undetectable activity (**Fig 6D**). Therefore, our linear DNA cell-free expression approach provides an easy-to-use method for screening of enzymatic activity from among a list of enzyme variants.

**Figure 6.**
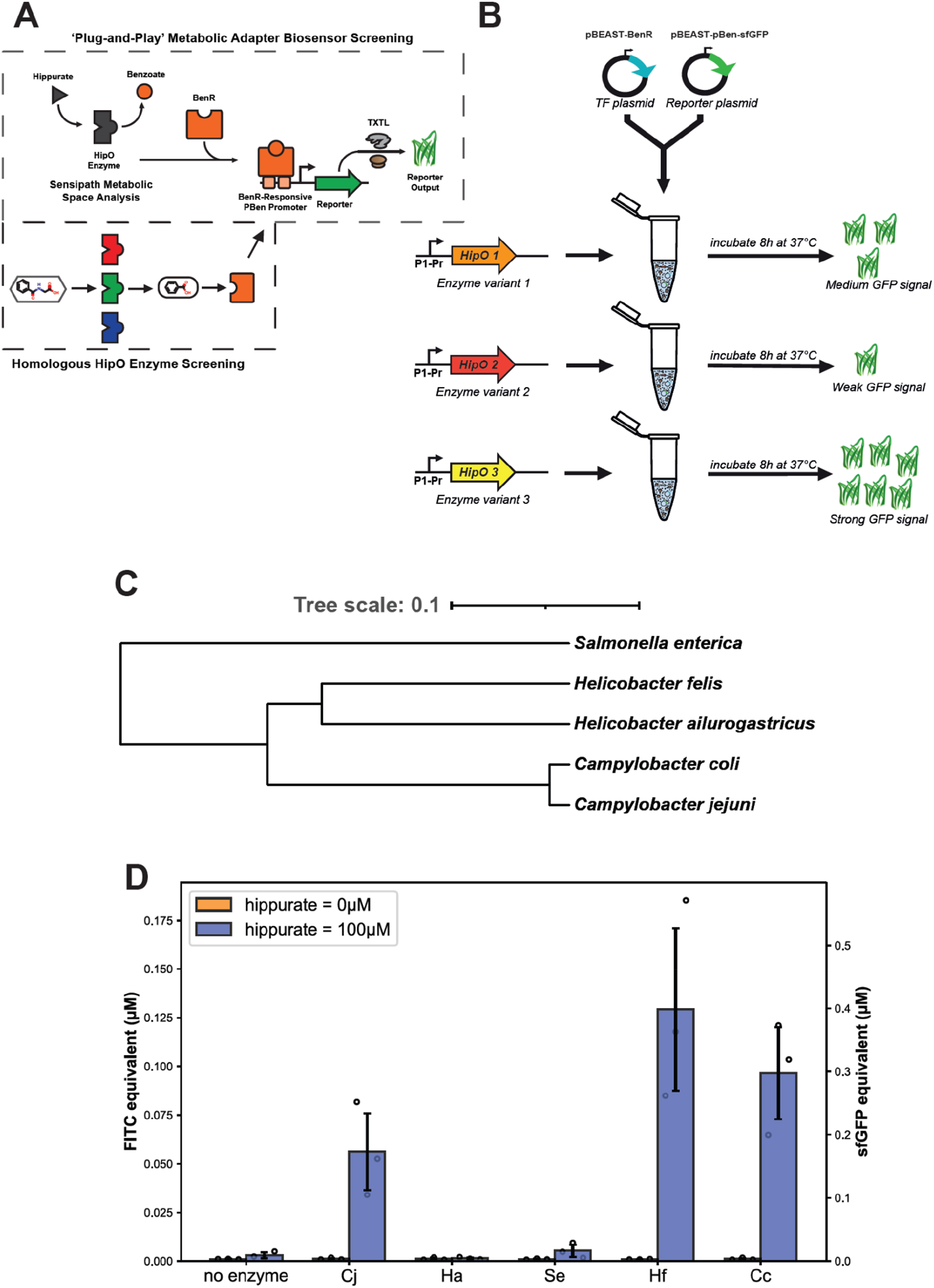
Δ*recBCD* extract enables enzyme variant screening from linear DNA template. **(A)** HipO enzyme variants are expressed in a cell-free reaction enabling conversion of hippurate into benzoate at various rates, acting as metabolic transducers. The produced benzoate triggers a genetically encoded biosensor by binding to the BenR transcription factor that activates the pBen promoter driving a GFP reporter protein. **(B)** Overview of the workflow used for the characterization of enzyme variants. Each tested sequence of enzyme is added as a linear DNA template to a different cell-free reaction containing the same pair of plasmids encoding the benzoate biosensor. The screening results are obtained after an 8 h incubation at 37 °C. **(C)** Phylogenetic tree showing the five different HipO homologs tested in this study. The tree was constructed following a methodology described in the method section. **(D)** Performance of HipO homologs expressed in cell-free from linear DNA templates. Endpoint fluorescence after 8 h of reaction shows the amount of reporter produced by the system for each enzyme candidate. Two of them (Hf & Cc) present a bioconversion efficiency at least as good as the original candidate (Cj), one (Se) demonstrates a weak conversion activity, and one variant (Ha) shows no activity. Data shown are mean±SD for 3 replicates done on different days. Additional experimental conditions are listed in Sup Table 6.

### Cell-free extract preparation and differential buffer calibration for resource-efficient prototyping: a workflow

In an attempt to provide the community with the most efficient method, we decided to comparatively evaluate the cost and time required to prototype a genetic part in one of the three systems where high expression efficiency is reached: plasmid DNA used in cell-free reaction, GamS protected linear DNA used in cell-free reaction, or linear DNA in cell-free extract of Δ*recBCD* strain origin (**Sup Fig 12**). The main differences observed were between the two linear DNA methods compared to the plasmid DNA method. Using linear DNA workflows reduces the time required to characterise a genetic part to a single day of experiments versus the traditional method that requires at least a week of work for similar results. The absence of cloning, plasmid verification by sequencing, and high-quantity DNA preparation steps in the linear workflows also reduces the cost of these methods by more than 50%. Finally, the use of Δ*recBCD* strain allows eliminating the cost of commercial GamS purchase from the final price, which comparatively reduces the cost of DNA part prototyping in this system. Taken together, these estimates argue for the use of the Δ*recBCD* strain based workflows for large scale DNA parts prototyping in cell-free systems. We established a robust and versatile workflow to unlock the potential of these optimized methods by non-specialists (**Fig 7**). It describes in parallel the preparation of the home made cell-free systems and the linear DNA required to test the desired genetic parts. The main differences with classical cell-free prototyping protocols are present in two parts of the workflow: the DNA design and build steps that only consist of PCR amplifying expressions cassettes and the extract buffer calibration step where the optimal magnesium glutamate and potassium glutamate concentrations have to be determined using a linear DNA template for reporter expression.

**Figure 7.**
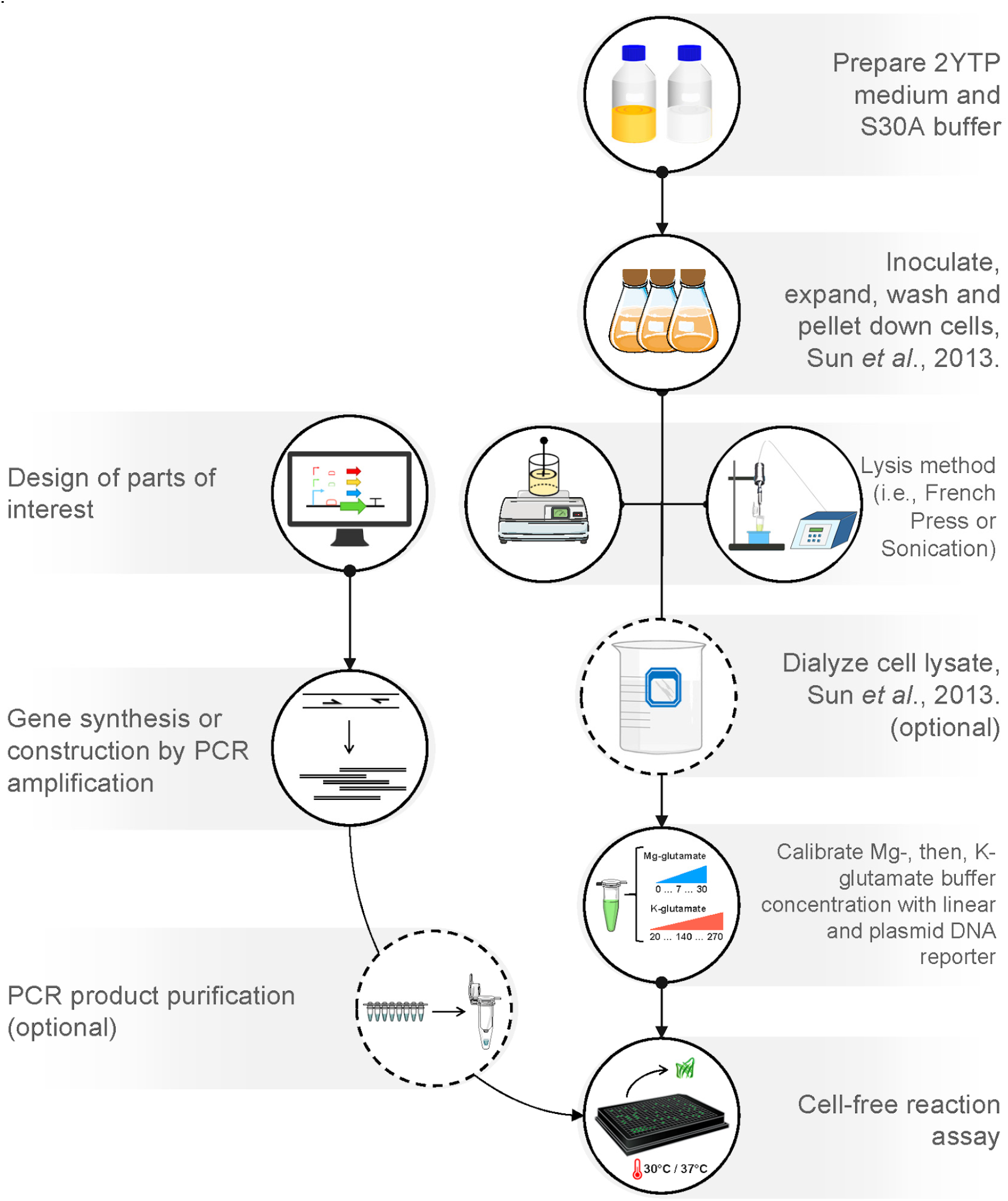
Workflow for linear DNA specific optimization. Linear DNA expression can be quickly characterized by using BL21 Δ*recBCD* strains developed in this work. Parts can be designed and synthesized, or generated by PCR assembly, and applied directly to the cell-free master mix. Independent of the lysis step (Sonication or French Press), in order to achieve optimal gene expression with linear or plasmid DNA, the buffer should be calibrated specifically for that DNA type. Once the optimal buffer composition for linear DNA templates has been identified, reactions using linear DNA can be efficiently performed.

## DISCUSSION

Here, we deleted the *recB* or *recBCD* genes from the *E. coli* BL21 Rosetta2 genome and showed that extracts prepared from these strains support robust protein expression from linear DNA templates without further supplementation of reagents or DNA modifications. Importantly, we demonstrated that specifically calibrating the cell-free buffer using linear DNA templates leads to higher expression from σ70 promoters in linear DNA, reaching near-plasmid levels for equimolar DNA concentrations. Interestingly, the buffer optimized for linear DNA expression had systematically lower K-glutamate levels. We found that the increased linear DNA expression at lower K-glutamate concentrations is not accompanied by increased DNA stability. These data suggest that high levels of K-glutamate could be inhibiting transcription from linear DNA, in agreement with previous reports ^41,42^. In all, our results highlight the importance of specifically tailoring the buffer optimization procedure for the type of DNA (linear or plasmid) to be used in the final application.

We showed that Δ*recBCD* extracts support rapid screening and characterization of genetic constructs for two relevant applications: toehold switch library characterization, and enzyme activity screening. While this manuscript was being prepared for submission, consistent with our findings a paper published by Arce *et al*. showed that cell-free extract depleted of the RecBCD exonuclease by CRISPRi can be used for prototyping of toehold switch biosensors encoded on linear DNA templates ^57^. Our method, by avoiding cloning and PCR purification, provided significant gains in time and cost. Synthetic linear DNA fragments could be ordered from a commercial provider and directly amplified and characterized the same day upon reception. The same process would have taken at least one week using plasmid DNA, if one takes into account the PCR, cloning, sequence verification, and maxiprep steps (**Sup Fig 12**). Given the current improvement in the speed of DNA synthesis and delivery, we estimate that a complete Design-Build-Test cycle can be performed in our linear cell-free set-up within five business days. Coupled with automated liquid handling technologies, such as acoustic droplet dispensing systems ^36,58,59^, linear DNA expression can further scale up the throughput of design exploration in cell-free applications substantially.

Our results are robust and repeatable, within and across laboratories. Δ*recBCD* extracts produced consistent results across biological replicates and different extract batches prepared using different strain backgrounds (-pRARE2 and +pRARE2). We also tested the robustness of the Δ*recBCD* extracts across different experimental parameters: different cell lysis methods (French press vs Sonication), temperatures (37 °C and 30 °C), DNA templates (*sfGFP, eGFP*), and different DNA concentrations (see **Sup Table 6** for details). While there were differences in GFP expression observed across different batches and experimental conditions, in two different laboratories (in Paris and Montpellier), the Δ*recBCD* extracts consistently supported strong protein expression (near-plasmid levels) from unprotected linear DNA templates after buffer calibration using linear DNA.

We anticipate that the strains and methods provided here will facilitate the use of linear DNA templates in cell-free systems and accelerate the DBT cycles for the synthetic biology community. Our data suggest that deletion of the exonuclease genes from the parental strain can be a general strategy for using linear DNA templates in a wide range of species of interest for cell-free synthetic biology, such as *Vibrio natriegens, Bacillus subtilis, Salmonella enterica*, and *Corynebacterium glutamicum* ^21^. The Δ*recB* and Δ*recBCD* BL21 strains are available to the scientific community through Addgene (**Sup Table 1**).

## Materials and Methods

### Bacterial strains and growth conditions

Strains BL21 Rosetta2 and BL21 DE3 were obtained from Merck Millipore, and the former was used to generate the genomic knockouts. Following the loss of the pRARE2 plasmid during the λ-red recombination protocol (-pRARE2 variants), all the three genotypes (WT, Δ*recB*, and Δ*recBCD*) were retransformed to generate both the +pRARE2 variants. All *E. coli* strains used for cell extract preparation in this work are listed in **Sup Table 1**. Further, *E. coli* KL740 cl857+ (Arbor Biosciences #502000) was used to maintain the P70a-deGFP plasmid for maxipreps (**Sup Table 4**), and *E. coli* strains TOP10 and DH5α were used for cloning. Cells were grown for maintenance and propagation in LB media (liquid, or solid with 1.5% w/v agar) at 37 °C and supplemented with the appropriate antibiotics depending on the plasmid(s) they carry (see **Sup Tables 1 and 4**) (Cam^R^: 10 µg mL^-1^ chloramphenicol, Amp^R^: 100 µg mL^-1^ ampicillin, Kan^R^: 25 µg mL^-1^ kanamycin, Spec^R^: 100 µg mL^-1^ spectinomycin). For extract preparation, cells were grown similarly, but in 2xYT-P media (described below).

### Genomic knockouts

*E. coli* BL21 Rosetta2 was used to make genomic deletions Δ*recB* and Δ*recBCD* using λ-Red recombination ^34^. Briefly, Rosetta2 was transformed with the helper plasmid pKD46 carrying the λ-Red recombination genes (γ, β, exo), followed by transformation with linear FRT-Kan^R^-FRT fragments, PCR amplified from the template plasmid pKD13 (**Fig 1A**). Two types of FRT-Kan^R^-FRT fragments were amplified using Platinum Polymerase II or Phusion polymerase (Thermo Fisher) with different flanking homology arms (41-43 bp) using primers P1 & P2 (for deleting *recB*) or primers P3 & P4 (for deleting *recC, ptrA, recB*, and *recD*). Kanamycin-resistant colonies were screened by colony PCR using primers P5 & P6 or primers P6 & P7, respectively (data not shown). After confirmation of recombination, cells were transformed with pCP20 plasmid carrying the thermo-sensitive flippase (*flp*) that was subsequently induced at 43 °C overnight for removal of *Kan*^*R*^ from the genome and curing of the pCP20 plasmid ^60^. Next, kanamycin-sensitive colonies were screened by colony PCR using primers P5 & P6 or primers P6 & P7 (**Fig S1**), and the genomic deletions verified by Sanger sequencing (**Fig 1B**). Sequences of all primers used are listed in **Sup Table 3**.

### Plasmid, linear, and Chi DNA

Plasmids P70a-deGFP and CMP (**Sup Table 4**) were used as expression templates for the deGFP and sfGFP fluorescent proteins, respectively. P70a-deGFP was maintained in *E. coli* KL740 cl857+ and CMP was maintained in *E. coli* DH5α cells for maxiprep using the NucleoBond Xtra Maxi kit (Macherey-Nagel). Linear DNA fragments used as expression templates in the cell-free reactions were PCR amplified using primers and templates listed in **Sup Table 7**. PCRs were performed using either: (i) the Q5 High-Fidelity 2x Master Mix (NEB), followed by DpnI (NEB) treatment and DNA purification with Monarch Plasmid Miniprep kit (NEB), or (ii) with Invitrogen Platinum SuperFi II and used without additional purification steps. In each case, PCR products were verified on a 1% agarose gel (1x TAE) prior to use. Purified DNA concentration was measured using a Nanodrop. However, non-purified PCR products were quantified via QuBit (Qubit dsDNA BR Assay Kit) since the Nanodrop overestimates the concentration of unpurified reactions by >20%, possibly due to ssDNA absorbance.

Chi6 forward and reverse oligonucleotides (P11 and P12 in **Sup Table 3**) were ordered from IDT and annealed according to Marshall et al. ^20^. The oligonucleotides were mixed together in equimolar concentrations (80 µM) in miliQ water. The strands were annealed by heating to 94°C for 2 minutes and gradually cooling to room temperature. The Chi6 hybridized oligonucleotides concentration was also measured using QuBit.

Plasmids sTR060 and sTR056 (**Sup Table 5**) were constructed using standard molecular cloning methods to express the gen2_num01 trigger and toehold-switch pair from Green et al. ^48^. All toehold switch variants for linear DNA expression were PCR amplified using two consecutive PCRs. The first PCR used to add the variant RBS was done using sTR056 plasmid as template, generating the twelve switch fragments prefixed ‘sw_’ (**Sup Table 7**). The second PCR was used to add the promoter P1 upstream of the toehold switch, generating the twelve transcription unit fragments prefixed ‘P1-’ (**Sup Table 7**) that were used in the cell-free reactions.

Plasmids pBEAST-BenR and pBEAST-pBen-sfGFP used as expression templates for the BenR transducer and benzoate-inducible sfGFP were maintained in *E. coli* DH5α for maxiprep (**Sup Table 4**) ^12^. HipO enzyme variants for linear DNA expression were PCR amplified using primers p33 and p34 (**Sup Table 3**) and five template gBlocks gene fragments ordered from IDT, suffixed ‘Gblocks’ in **Sup Table 5**.

### Cell-free extract preparation

Cell-free extracts were generated from both WT and knockout BL21 strains by physical disruption, either by sonication or French press lysis (**Sup Table 2**). The extract preparation protocol adapted from Sun et al. ^31^ is briefly presented here. Cells were inoculated from an agar plate into 2xYT-P media and grown overnight at 37 °C. The next day, they were subcultured into fresh 2xYT-P and grown to an OD_600_ of 1.5-2.0. For cells containing the pRARE2 and/or the pBADmod1-linker2-gamS plasmid (**Sup Table 1**), appropriate antibiotics were added to the media. For cell-free extracts with overexpressed GamS (**Sup Table 2, Sup Fig 8**), cells were grown in 2xYT-P with ampicillin and induced at an OD_600_ of 0.4-0.6 with 0.25% w/v L-arabinose, after which they were grown at 30 °C until they reached an OD_600_ of 1.5 to 2.0. Cells were spun down and washed in 200 mL S30A buffer twice. Finally, they were spun down twice into a compact pellet, the supernatant fully removed, and the pellet frozen at -80 °C. On day 3, the pellet was resuspended in 0.9-1 mL of S30A buffer per gram of its weight. For the **sonication lysis** method, 1 mL each of the S30A resuspended cells were aliquoted into 1.5 mL microtubes and lysed on ice using Vibra-Cell CV334 sonicator (3 mm probe), with a setup of 20% amplitude for 3 cycles (30 s sonication, 1 min pause). Next, the extract was pelleted down at 12000xg for 10 min at 4 °C and the supernatant was incubated at 37 °C for 80 minutes with 200 rpm agitation. Finally, supernatant was centrifuged again at 12000xg for 10 min at 4 °C, and the pellet-free cell extract was flash frozen and stored at -80 °C prior to buffer calibration. For the **French press lysis** method, the Avestin EmulsiFlex-C3 homogeniser was used at 15000-20000 psi and the final pellet-free cell extract was dialysed (using a Spectra/Por 4 Dialysis Tubing, 12-14 kD MWCO from Repligen) overnight in S30B buffer before one final centrifugation at 12000xg for 30 minutes, aliquoted, and flash frozen.

### Cell-free buffer calibration

Cell-free buffer was calibrated for optimal magnesium glutamate and potassium glutamate concentrations as described in Sun et al. ^31^. First, Mg-glutamate was calibrated with a fixed K-glutamate concentration (80 mM). Next, K-glutamate was calibrated using the optimal Mg-glutamate concentration identified for the highest GFP expression. Each extract was calibrated with 1 nM or 4 nM of linear or plasmid DNA with Mg-glutamate/K-glutamate concentrations described in **Figs 2B** and **2C, Sup Figs 5B** and **5C**, or **Sup Figs 8B** and **8C**. The endpoint fluorescence measurements were taken after 8 h at 37 °C or 30 °C (details in **Sup Table 6**).

### GamS expression and purification

The λ-phage GamS protein (with N-terminal 6xHis tag) was purified as described previously ^19^, and used at an optimal concentration of 2 μM in our cell-free reactions. The GamS expression plasmid (pBADmod1-linker2-gamS) was ordered from Addgene (#45833). GamS expression was achieved in a culture of 4x 400 mL LB grown at 37°C until an OD_600_ of ∼0.6, followed by induction with 0.2% w/v arabinose and incubation at 37°C for 3 h. The cells were harvested by centrifugation at 6000x rpm for 20 min and the pellets stored overnight at -40 °C. The pellets were resuspended (30 mL per pellet) in a lysis buffer (Tris 50 mM, NaCl 150 mM, DTT 2 mM, pH 8), followed by addition of lysozyme, sonication, and centrifugation at 18000x rpm for 30 min. GamS was purified using Histrap columns, with an elution buffer (Tris 50 mM, NaCl 150 mM, DTT 2 mM, Imidazole 300 mM, pH 8). After the column purification, the protein samples were pooled, dialyzed overnight at 4 °C (Tris 50 mM, NaCl 150 mM, DTT 2 mM, pH 8), and concentrated to OD_280_ 8.9 using a Centricon 3 kDa filter. To ensure maximal purity, gel filtration using a HiLoad 16/60 Superdex 75 column from Sigma-Aldrich was conducted and the fractions containing the protein were pooled and concentrated at OD_280_ 0.6453, equivalent to 1.5 mg/ml, before being aliquoted and stored at -20°C. These fractions were confirmed on SDS-PAGE.

### RNA production and purification

Trigger RNA gen2_num01 was produced from the sTR060 DNA template after PCR-amplification using primers 3116 and 2863 (**Sup Table 3**), with a T7 promoter upstream of the trigger sequence. *In vitro* transcription of the trigger RNA was performed using the TranscriptAid T7 High Yield Transcription Kit (Thermo Scientific), following the manufacturer’s protocol. After transcription, the DNA template was removed by digestion with the kit-supplied RNase-free DNase I enzyme and the product was purified using Monarch RNA Cleanup Kit (50 μg). The concentration of the final product was determined by A260 measurement using a Nanodrop spectrometer and the RNA was diluted to 40 µM (20X working concentration) before being aliquoted and stored at -80°C.

### Search for enzyme variants

The previously characterized *Campylobacter jejuni* HipO enzyme ^12^ was used to identify four additional enzyme variants from the UniRef90 reference cluster (https://www.uniprot.org/uniref/UniRef90_P45493), which contains members with at least 90% identity to the original sequence (UniProt P45493) ^61^. The 4 homologs chosen are representative members of the four biggest clusters that group together more than 93% of all the existing UniProt entries homologous to the original sequence. The phylogenetic tree was generated from the amino acids sequences using the Clustal Omega multiple sequence alignment tool (https://www.ebi.ac.uk/Tools/msa/clustalo/). The resulting tree was then formatted by iTOL (https://itol.embl.de/).

### Fluorescence measurements

Quantitative measurements of gene expression were carried out using the reporter proteins deGFP and sfGFP. Fluorescence was monitored with a Biotek Synergy H1M reader, a Cytation 3 Reader, or a Synergy HTX plate reader (Biotek Instruments, Ex 485 nm, Em 528 nm). 384-well square- or round-bottom microplates were used (**Sup Table 6**) and covered by an adhesive plate seal (Thermo Scientific #AB0558 or #232701) to minimize evaporation. Kinetic runs recorded fluorescence data at regular intervals for 8 hours (**Sup Fig 3**). End-point measurements were reported after 8 h of incubation at either 30 °C or 37 °C. To be able to compare our fluorescence data between different plate readers and laboratories, raw data collected was converted to FITC Mean Equivalent Fluorescence (MEF) values for 20 µL reactions at 30 °C or 37 °C according to Jung et al. ^18^. FITC MEF values were converted to equivalent fluorescent protein concentrations using calibration curves made using recombinant eGFP from ChromoTek GmbH (reference EGFP-250) and sfGFP purified in-house (kind gift from Team SyBER at the Micalis Institute).

### qPCR on cell-free DNA

To investigate DNA stability in different K-glutamate concentrations, linear or plasmid DNA was incubated in low K-glutamate (optimal for linear DNA) or high K-glutamate (optimal for plasmid DNA) buffer compositions in 10.5 µL cell-free reactions. BL21_7, ΔrecB_8 and ΔrecBCD_10 extracts (**Sup Table 2**) were incubated with 1 nM DNA at 30°C for 3 different durations (0 h, 1 h, and 8h). As described above, fluorescence from each reaction was acquired using a plate reader. At each time-point, 8 µL of the cell-free reaction was taken out, mixed with EDTA (end-concentration 11 mM), and boiled at 70 °C for 30 minutes to stop the exonuclease activity. Next, DNA was purified with Monarch PCR & DNA Cleanup Kit (NEB) and eluted in 20 µL of nuclease-free water. Standard curves for the qPCR were prepared with serial dilutions of known linear DNA concentrations. The qPCR reactions to quantify *deGFP* used 2 µM each of primers P13 & P14 and 1 µM of probe P15 ordered from Eurofins Genomics (**Sup Table 3**), as previously reported ^62^. qPCR reactions (10 µL each) were performed with OneTaq polymerase (NEB #M0484) in 96 well PCR plate (4titude 4ti-0750), with a clear qPCR seal (4titude 4Ti-0560), and incubated in a thermal cycler (Eppendorf Realplex 2 Mastercycler) with the following program: 95 °C for 3 min; 40x cycles of (94 °C for 15 sec, 53 °C for 30 sec, 68°C for 30 sec).

## Author Contributions

ACB, AL, CLB, JB, MK conceived the study. ACB, AL, PS, PLV, MK designed the experiments. ACB, ADV, AL, MCG, PS, PLV, TA, TR-T performed the experiments. ACB, AL, JB, PS, PLV, MK analysed the results. CLB, JB, JLF, MK acquired the funding. JB, MK wrote the manuscript with contributions from all authors. All authors read and approved the final manuscript.

The authors declare no competing financial interest.

## Supporting Information

1. Supplementary Information file (PDF).
2. Supplementary Figures file (PDF).

## Acknowledgements

We thank Mahnaz Sabeti Azad for her comments on the manuscript, and Stephen McGovern (Team SyBER, Micalis Institute) for the kind gift of the purified sfGFP protein. ACB and JLF acknowledge funding support by the ANR SINAPUV grant (ANR-17-CE07-0046). PS, TRT, JLF and JB acknowledge funding support by the ANR SynBioDiag grant (ANR-18-CE33-0015). JLF acknowledges funding support by the ANR iCFree grant (ANR-20-BiopNSE). JB acknowledges support from the ERC starting “COMPUCELL” (grant number 657579). The Centre de Biochimie Structurale acknowledges support from the French Infrastructure for Integrated Structural Biology (FRISBI) (ANR-10-INSB-05-01). MK acknowledges funding support from INRAe’s MICA department, Université Paris-Saclay, Ile-de-France (IdF) region’s DIM-RFSI, and ANR DREAMY (ANR-21-CE48-003). This work was supported by funding through the European Research Council Consolidator Award (865973 to CLB).

