## Supplementary Information file for "Differentially optimized cell-free buffer enables robust expression from unprotected linear DNA in exonuclease-deficient extracts"

**Supplementary Table 1**

*Escherichia coli* strains used for cell extracts made in the study:

| Strain | Genotype | Plasmid | Source |
|---|---|---|---|
| BL21 DE3 | F- *ompT hsdS*B(rB- mB-) *gal dcm* (DE3) | - | Merck Millipore #70954 |
| BL21 | F- *ompT hsdS*B(rB- mB-) *gal dcm* | - | Merck Millipore #71402, after loss of the pRARE2 plasmid |
| BL21 Δ*recB* | F- *ompT hsdS*B(rB- mB-) *gal dcm* Δ*recB* | - | This work (Addgene #176580) |
| BL21 Δ*recBCD* | F- *ompT hsdS*B(rB- mB-) *gal dcm* Δ(*recC-recD)* | - | This work (Addgene #176581) |
| BL21 Rosetta2 | F- *ompT hsdS*B(rB- mB-) *gal dcm* | pRARE2 ($Cam^R$) | Merck Millipore #71402 |
| BL21 Rosetta2 Δ*recB* | F- *ompT hsdS*B(rB- mB-) *gal dcm* Δ*recB* | pRARE2 ($Cam^R$) | This work (Addgene #176582) |
| BL21 Rosetta2 Δ*recBCD* | F- *ompT hsdS*B(rB- mB-) *gal dcm* Δ(*recC-recD)* | pRARE2 ($Cam^R$) | This work (Addgene #176583) |
| BL21 Rosetta2 GamS | F- *ompT hsdS*B(rB- mB-) *gal dcm* | pRARE2 ($Cam^R$), pBADmod1-linker2-gamS ($Amp^R$) | This work (Addgene #176584) |
| BL21 Rosetta2 Δ*recB* GamS | F- *ompT hsdS*B(rB- mB-) *gal dcm* Δ*recB* | pRARE2 ($Cam^R$), pBADmod1-linker2-gamS ($Amp^R$) | This work (Addgene #176585) |
| BL21 Rosetta2 Δ*recBCD* GamS | F- *ompT hsdS*B(rB- mB-) *gal dcm* Δ(*recC-recD)* | pRARE2 ($Cam^R$), pBADmod1-linker2-gamS ($Amp^R$) | This work (Addgene #176586) |

**Supplementary Table 2**

Description of the different extracts used in the study:

| **Extract** | **Strain*** | **Lysis method** |
|---|---|---|
| BL21_1 | BL21 Rosetta2 | Sonication |
| Δ*recB*_2 | BL21 Δ*recB* | Sonication |
| Δ*recBCD*_3 | BL21 Δ*recBCD* | Sonication |
| BL21_4 | BL21 DE3 | French Press |
| Δ*recBCD*_5 | BL21 Δ*recBCD* | French Press |
| Δ*recBCD*_6 | BL21 Δ*recBCD* | French Press |
| BL21_7 | BL21 Rosetta2 | Sonication |
| Δ*recB*_8 | BL21 Rosetta2 Δ*recB* | Sonication |
| Δ*recBCD*_9 | BL21 Rosetta2 Δ*recBCD* | Sonication |
| Δ*recBCD_10* | BL21 Rosetta2 Δ*recBCD* | Sonication |
| BL21_11 | BL21 Rosetta2 GamS | Sonication |
| Δ*recB*_12 | BL21 Rosetta2 Δ*recB* GamS | Sonication |
| Δ*recBCD*_13 | BL21 Rosetta2 Δ*recBCD* GamS | Sonication |

*As used in Supplementary Table 1

**Supplementary Table 3**
All primers used in this study:

| Primer | Sequence (5'->3') |
|---|---|
| ***ΔrecB/ΔrecBCD* knockouts** | |
| P1 (recB_KO_F) | tgctccacagcttccagtaattgcttttgcaatttcattaattcc ggggatccgtcgacc |
| P2 (recB_KO_R) | cagcaaacaatgcccctgatgagtgaaaagaatgagtg atgtgtaggctggagctgcttc |
| P3 (recD_KO_F) | catgcgtaacactcgtgcgtcgcatccggcaattacgttaa ttccggggatccgtcgacc |
| P4 (recC_KO_R) | cgcctgcattgcccgaatcgtcagtagtcaggagccgcttg tgtaggctggagctgcttc |
| P5 (recB_screen_F) | tattttccagtcgtgaaagc |
| P6 (recB_screen_R) | ttgctgatttcttccatcag |
| P7 (recD_screen_F) | ttgatttactgcccgagagc |
| P8 (recC_screen_R) | gtcaaccgaatgcagacatc |
| P9 (VF2) | tgccacctgacgtctaagaa |
| P10 (VR) | attaccgcctttgagtgagc |
| ***qPCR*** | |
| P13 (EGFP-1-F) | GACCACTACCAGCAGAACAC |
| P14 (EGFP-2-R) | GAACTCCAGCAGGACCATG |
| P15 (EGFP-FAM-BBQ650) | [FAM]-AGCACCCAGTCCGCCCTGAGCA-[BBQ650] |
| ***Chi DNA*** | |
| P11 (chi6.fwd) | TCACTTCACTGCTGGTGGCCACTGCTGG TGGCCACTGCTGGTGGCCACTGCTGGT GGCCACTGCTGGTGGCCACTGCTGGTG GCC |
| P12 (chi6.rev) | GGCCACCAGCAGTGGCCACCAGCAGTG GCCACCAGCAGTGGCCACCAGCAGTGG CCACCAGCAGTGGCCACCAGCAGTGAA GTGA |
| ***sfGFP reporter & HipO enzymes' amplification*** | |

| | |
|---|---|
| p33 | CTCGGATACCCTTACTCTGTTGAAAAC |
| p34 | TCTTACTGAAGCACGATTCTACTCGG |
| ***Toehold switches' amplification*** | |
| toehold_P1_add_FW | gggtgagctaacaccgtgcgtgttgacaattttacctctggcggtgataatggttgcaagcaGGGTCTTATCTTATCTATCTCGTTTATCCCTG |
| J61100_FW | GGGTCTTATCTTATCTATCTCGTTTATCCCTGCAtacagaaagaggggacatATGCAATGATAAACGAGAACCTGG |
| J61101_FW | GGGTCTTATCTTATCTATCTCGTTTATCCCTGCAtacagaaagacaggacctATGCAATGATAAACGAGAACCTGG |
| J61105_FW | GGGTCTTATCTTATCTATCTCGTTTATCCCTGCAtacagaaagacatgacgtATGCAATGATAAACGAGAACCTGG |
| J61106_FW | GGGTCTTATCTTATCTATCTCGTTTATCCCTGCAtacagaaagataggagatATGCAATGATAAACGAGAACCTGG |
| J61102_FW | GGGTCTTATCTTATCTATCTCGTTTATCCCTGCAtacagaaagatccgatgtATGCAATGATAAACGAGAACCTGG |
| J61103_FW | GGGTCTTATCTTATCTATCTCGTTTATCCCTGCAtacagaaagattagacatATGCAATGATAAACGAGAACCTGG |
| J61104_FW | GGGTCTTATCTTATCTATCTCGTTTATCCCTGCAtacagaaagaagggacatATGCAATGATAAACGAGAACCTGG |
| J61109_FW | GGGTCTTATCTTATCTATCTCGTTTATCCCTGCAtacagaaagactggagatATGCAATGATAAACGAGAACCTGG |
| J61107_FW | GGGTCTTATCTTATCTATCTCGTTTATCCCTGCAtacagaaagaagagacttATGCAATGATAAACGAGAACCTGG |
| J61108_FW | GGGTCTTATCTTATCTATCTCGTTTATCCCTGCAtacagaaagacgagatatATGCAATGATAAACGAGAACCTGG |
| J61110_FW | GGGTCTTATCTTATCTATCTCGTTTATCCCTGCAtacagaaagaggcgaattATGCAATGATAAACGAGAACCTGG |
| SBa_000587_RV | acaaaaaacccctagccgcc |

| | |
|---|---|
| ***DNA template amplification for Trigger RNA transcription*** | |
| 3116 | CCCGCGAAATTAATACGACTC |
| 2863 | atccggatatagttcctcct |
| ***Modified DNA-ends' amplification*** | |
| SP0_3 2-MethoxyEtoxy | /52MOErC/ctcggatacccttactctgttgaaaacgaatagataggtt |
| SPN_3 2-MethoxyEtoxy | /52MOErT/tcttactgaagcacgattctactcggaagtggtcaataat |
| SP0_1 Phosphorothioate | ctc*g*g*atacccttactctgttgaaaacgaatagataggtt |
| SPN_1 Phosphorothioate | tct*t*a*ctgaagcacgattctactcggaagtggtcaataat |
| SP0_5 FluoroC | /52FC/ctcggatacccttactctgttgaaaacgaatagataggtt |
| SPN_5 FluoroC | /52FC/tcttactgaagcacgattctactcggaagtggtcaataat |
| SP0_2 Phosphorylation | /5Phos/ctcggatacccttactctgttgaaaacgaatagataggtt |
| SPN_2 Phosphorylation | /5Phos/tcttactgaagcacgattctactcggaagtggtcaataat |

**Supplementary Table 4**
Plasmids used in this study:

| Plasmid | Antibiotic Selection | Accession/ Catalog reference | Source |
|---|---|---|---|
| P70a-deGFP | $Amp^R$ | Arbor# 502056 | Arbor Biosciences |
| pKD46 | $Amp^R$ | NCBI# AY048746.1 | (Datsenko and Wanner 2000) |
| pKD13 | $Amp^R$ / $Kan^R$ | NCBI# AY048744.1 | (Datsenko and Wanner 2000) |
| pCP20 | $Amp^R$ / $Cam^R$ | CGSC# 64621 | (Cherepanov and Wackernagel 1995) |
| pBADmod1-linker2-gamS | $Amp^R$ | Addgene #45833 | - |
| pBEAST-BenR | $Amp^R$ | Addgene #114597 | (Voyvodic et al. 2019) |
| pBEAST-pBen-sfGFP | $Amp^R$ | Addgene #114598 | (Voyvodic et al. 2019) |
| CMP | $Kan^R$ | Addgene #177368 | This work |
| sTR056 | $Spec^R$ | Addgene #177369 | This work |
| sTR060 | $Amp^R$ | Addgene #177370 | This work |

**Supplementary Table 5**

DNA sequences for constructs used in the study:

| Plasmid/ DNA construct | Sequence (5’->3’) |
|---|---|
| CMP<br><br>[Pr7- RiboJ-BCD2<br>-sfGFP] | aaaaaatttatttgctttcgcatctttttgtacctataatgtgtggaggatccagctgtcaccggatgtgct<br>ttccggtctgatgagtccgtgaggacgaaacagcctctacaaataattttgtttaagggcccaagttc<br>acttaaaaaggagatcaacaatgaaagcaattttcgtactgaaacatcttaatcatgctaaggaggt<br>tttctaatgtcaaaaggagaagaactttttacaggtgtagtacctatcttggttgaattggatggtgatgt<br>taacggtcacaaattttctgtacgtggtgaaggtgaaggtgatgcaactaacggtaaattgacactta<br>aattcatttgtacaactggaaaacttcctgttccttggcctactcttgttacaacattgacatatggagta<br>caatgtttttcacgttatcctgatcatatgaaacgtcacgatttttttaaatctgctatgccagaaggttat<br>gtacaagaacgtacaatttcatttaaagatgacggaacatataaaacacgtgctgaagtaaaattc<br>gaaggtgacactcttgttaatcgtatcgaattgaaaggaatcgatttcaaagaagatggtaacatttt<br>gggacacaaacttgaatacaacttcaactctcataatgtttatatcacagctgacaaacaaaaaaa<br>cggtattaaagctaattttaaaattcgtcacaatgttgaagatggatctgttcaattggctgatcattatc<br>aacaaaatacaccaatcggagacggaccagtattgcttccagataaccactacctttctactcaatc<br>agttctttcaaaagatcctaacgaaaaacgtgaccatatggtacttcttgaatttgttacagcagcag<br>gtatcactcacggtatggacgaactttataaataaactttatctgagaatagtcaatcttcggaaatcc<br>caggtggcatgctaaaagtctcgtaaagcgttctatcaataacccgttggtgccaggcatcaaataa<br>aacgaaaggctcagtcgaaagactgggcctttcgttttatctgttgtttgtcggtgaacgctctctacta<br>gagtcacactggctcaccttcgggtgggcctttctgcgtttataccgtctcagaatcggccgtgaaca<br>ataaaatagtttcggtattattgaccacttccgagtagaatcgtgcttcagtaagagagtcactaagg<br>gttagttagttagattagcagaaagtcaaaagcctccgaccggaggcttttgactaaaacttcccttg<br>gggttatcattggggctcactcaaaggcggtaatcagataaaaaaaatccttagctttcgctaagga<br>tgatttctgctagtattattagaaaaactcatcgagcatcaaatgaaactgcaatttattcatatcagga<br>ttatcaataccatatttttgaaaaagccgtttctgtaatgaaggagaaaactcaccgaggcagttcca<br>aagaatggcaaggtcctggtaacggtctgcgattccgacccgtccaacatcaatacaacctattaa<br>tttcccctcgtcaaaaataaggttatcaagtgagaaatcaccatgagtgacgactgaatccggtga<br>gaatggcaagagcttgtgcatttctttccagacttgttcaacaggccagccattacgctcgtcatcaa<br>aatcactcgcatcaaccaaaccgttattcatgcgtgattgcgcctgagcaagacgaaatacacgat<br>cgctgttaaaaggacaattacaaacaggaatcgaatgtaaccggcgcaggaacacggccagcg<br>catcaacaatattttcacctgaatcaggatattcttctaatacctggaaggctgttttcccaggaatcgc<br>ggtggtgagtaaccacgcatcatcaggagtacggataaaatgcttgatggtcgggagaggcataa<br>actccgtcagccagttgagacggaccatctcatctgtaacatcattggcaacgctacctttgccatgtt<br>tcagaaacaactctggcgcatcgggcttcccatacaagcgatagattgtcgcacctgattgcccga<br>cattatcgcgagcccatttatacccatataaatcagcgtccatgttggagtttaagcgcggacggga<br>gcaagacgtttcccgttgaatatggctcataacaccccttgtattactgtttatgtaagcagacagtttta<br>ttgttcatgatgatatatttttatcttgtgcaatgtaacatcagagattttgagacacaacgtggctttgttg<br>aataaatcgaacttttgctgagttgaaggatcagctctagtagttacattgtcgatctgttcatggtgaa<br>cagctttaaatgcaccaaaaactcgtaaaagctctgatgtatctatcttttttacaccgttttcatctgtgc<br>atatggacagttttccctttgatatctaacggtgaacagttgttctacttttgtttgttagtcttgatgcttcac<br>tgatagatacaagagccataagaacctcagatccttccgtatttagccagtatgttctctagtgtggttc<br>gttgtttttgcgtgagccatgagaacgaaccattgagatcatgcttactttgcatgtcactcaaaaatttt<br>gcctcaaaactggtgagctgaatttttgcagttaaagcatcgtgtagtgtttttcttagtccgttacgtag<br>gtaggaatctgatgtaatggttgttggtattttgtcaccattcatttttatctggttgttctcaagttcggttac<br>gagatccatttgtctatctagttcaacttggaaaatcaacgtatcagtcgggcggcctcgcttatcaac<br>caccaatttcatattgctgtaagtgtttaaatctttacttattggtttcaaaacccattggttaagccttttaa<br>actcatggtagttattttcaagcattaacatgaacttaaattcatcaaggctaatctctatatttgccttgt<br>gagttttcttttgtgttagttcttttaataaccactcataaatcctcatagagtatttgttttcaaaagacttaa<br>catgttccagattatattttatgaattttttttaactggaaaagataaggcaatatctcttcactaaaaacta<br>attctaatttttcgcttgagaacttggcatagtttgtccactggaaaatctcaaagcctttaaccaaagg<br>attcctgatttccacagttctcgtcatcagctctctggttgctttagctaatacaccataagcattttcccta<br>ctgatgttcatcatctgagcgtattggttataagtgaacgataccgtccgttctttccttgtagggttttcaa |

| | |
|---|---|
| | tcgtggggttgagtagtgccacacagcataaaattagcttggtttcatgctccgttaagtcatagcgac<br>taatcgctagttcatttgctttgaaaacaactaattcagacatacatctcaattggtctaggtgattttaat<br>cactataccaattgagatgggctagtcaatgataattacatgtccttttcctttgagttgtgggtatctgta<br>aattctgctagacctttgctggaaaacttgtaaattctgctagaccctctgtaaattccgctagacctttg<br>tgtgttttttttgtttatattcaagtggttataatttatagaataaagaaagaataaaaaaagataaaaag<br>aatagatcccagccctgtgtataactcactactttagtcagttccgcagtattacaaaaggatgtcgc<br>aaacgctgtttgctcctctacaaaacagaccttaaaaccctaaaggcttaagtagcaccctcgcaa<br>gctcgggcaaatcgctgaatattccttttgtctccgaccatcaggcacctgagtcgctgtctttttcgtga<br>cattcagttcgctgcgctcacggctctggcagtgaatgggggtaaatggcactacaggcgccttttat<br>ggattcatgcaaggaaactacccataatacaagaaaagcccgtcacgggcttctcagggcgtttta<br>tggcgggtctgctatgtggtgctatctgactttttgctgttcagcagttcctgccctctgattttccagtctg<br>accacttcggattatcccgtgacaggtcattcagactggctaatgcacccagtaaggcagcggtatc<br>atcaacaggcttacccgtcttactgtccctagtgcttggattctcaccaataaaaaacgcccggcgg<br>caaccgagcgttctgaacaaatccagatggagttctgaggtcattactggatctatcaacaggagtc<br>caagcgagctcgtaaacttggtctgacagctctagctccggcaaaaaaacgggcaaggtgtcac<br>caccctgccctttttctttaaaaccgaaaagattacttcgcgtttgccacctgacgtctaagaaaagga<br>atattcagcaatttgcccgtgccgaagaaaggcccacccgtgaaggtgagccagtgagttgattgc<br>tacgtaattagttagttagcccttagtgactcctcggatacccttactctgttgaaaacgaatagatagg<br>ttgctagc |
| sTR056<br><br>[PrT7- trigger binding region- loop with RBS- linker-sfGFP] | taatacgactcactataggagcaGGGTCTTATCTTATCTATCTCGTTTATCCCT<br>GCATACAGAAACAGAGGAGATATGCAATGATAAACGAGAACCTGG<br>CGGCAGCGCAAAAGatgcGTAAAGGCGAAGAACTGTTTACCGGTGT<br>GGTTCCGATTCTGGTGGAACTGGACGGCGATGTTAATGGTCATAAA<br>TTCAGTGTTCGCGGCGAAGGTGAAGGCGATGCGACGAACGGCAAA<br>CTGACCCTGAAATTTATCTGCACCACGGGTAAACTGCCGGTCCCGT<br>GGCCGACGCTGGTGACCACGCTGACCTATGGCGTTCAATGTTTTG<br>CGCGTTACCCGGATCACATGAAACAGCACGACTTTTTCAAATCGGC<br>CATGCCGGAAGGCTATGTGCAGGAACGTACGATTAGCTTTAAAGA<br>CGATGGTACGTATAAAACCCGCGCGGAAGTGAAATTCGAAGGCGA<br>TACCCTGGTTAACCGTATCGAACTGAAAGGTATCGATTTCAAAGAA<br>GACGGCAATATTCTGGGTCATAAACTGGAATATAACTTCAATTCCC<br>ACAACGTGTACATCACCGCGGATAAACAGAAAAACGGCATTAAAGC<br>CAATTTCAAAATCCGCCATAATGTGGAAGATGGTAGCGTTCAGCTG<br>GCCGACCACTATCAGCAAAACACGCCGATTGGTGATGGCCCGGTC<br>CTGCTGCCGGACAATCACTACCTGAGTACCCAGTCCGTGCTGTCA<br>AAAGATCCGAACGAAAAACGTGACCACATGGTCCTGCTGGAATTTG<br>TGACGGCTGCGGGTATCACCCACGGCATGGACGAACTGTATAAAA<br>GGCCTgcagcaaacgacgaaaactacgctgcatcagttTAATAAtactcgaacccctagc<br>ccgctcttatcgggcggctaggggttttttgtCCTAGGGCGGCCGCGTCGTGACTG<br>GGAAAACCCTGGCGACTAGTCTTGGACTCCTGTTGATAGATCCAGT<br>AATGACCTCAGAACTCCATCTGGATTTGTTCAGAACGCTCGGTTGC<br>CGCCGGGCGTTTTTTATTGGTGAGAATCCAGGGGTCCCCAATAATT<br>ACGATTTACGTATTTAAATgaaccttgaccgaacgcagcggtggtaacggcgcagtg<br>gcggttttcatggcttgttatgactgttttttggggtacagtctatgcctcgggcatccaagcagcaagc<br>gcgttacgccgtgggtcgatgtttgatgttatggagcagcaacgatgttacgcagcagggcagtcg<br>ccctaaaacaaagttaaacatcATGAGGGAAGCGGTGATCGCCGAAGTATC<br>GACTCAACTATCAGAGGTAGTTGGCGTCATCGAGCGCCATCTCGA<br>ACCGACGTTGCTGGCCGTACATTTGTACGGCTCCGCAGTGGATGG<br>CGGCCTGAAGCCACACAGTGATATTGATTTGCTGGTTACGGTGAC<br>CGTAAGGCTTGATGAAACAACGCGGCGAGCTTTGATCAACGACCT<br>TTTGGAAACTTCGGCTTCCCCTGGAGAGAGCGAGATTCTCCGCGC<br>TGTAGAAGTCACCATTGTTGTGCACGACGACATCATTCCGTGGCGT<br>TATCCAGCTAAGCGCGAACTGCAATTTGGAGAATGGCAGCGCAAT |

| | |
|---|---|
| | GACATTCTTGCAGGTATCTTCGAGCCAGCCACGATCGACATTGATC<br>TGGCTATCTTGCTGACAAAAGCAAGAGAACATAGCGTTGCCTTGGT<br>AGGTCCAGCGGCGGAGGAACTCTTTGATCCGGTTCCTGAACAGGA<br>TCTATTTGAGGCGCTAAATGAAACCTTAACGCTATGGAACTCGCCG<br>CCCGACTGGGCTGGCGATGAGCGAAATGTAGTGCTTACGTTGTCC<br>CGCATTTGGTACAGCGCAGTAACCGGCAAAATCGCGCCGAAGGAT<br>GTCGCTGCCGACTGGGCAATGGAGCGCCTGCCGGCCCAGTATCA<br>GCCCGTCATACTTGAAGCTAGACAGGCTTATCTTGGACAAGAAGAA<br>GATCGCTTGGCCTCGCGCGCAGATCAGTTGGAAGAATTTGTCCAC<br>TACGTGAAAGGCGAGATCACCAAGGTAGTCGGCAAATAAGACaaTT<br>GTCCTTTTCCGCTGCATAACCCTGCTTCGGGGTCATTATAGCGATT<br>TTTTCGGTATATCCATCCTTTTTCGCACGATATACAGGATTTTGCCA<br>AAGGGTTCGTGTAGACTTTCCTTGGTGTATCCAACGGCGTCAGCC<br>GGGCAGGATAGGTGAAGTAGGCCCACCCGCGAGCGGGTGTTCCT<br>TCTTCACTGTCCCTTATTCGCACCTGGCGGTGCTCAACGGGAATCC<br>TGCTCTGCGAGGCTGGCCGTAGGCCGGCCtggtgaacagcttgaatgcacc<br>aaaaactcgtaaaagctctgatgtatctatctttttacaccgttttcatctgtgcatatggacagttttccc<br>tttgatatgtaacggtgaacagttgttctacttttgtttgttagtcttgatgcttcactgatagatacaagag<br>ccataagaacctcagatccttccgtatttagccagtatgttctctagtgtggttcgttgtttttgcgtgagc<br>catgagaacgaaccattgagatcatacttactttgcatgtcactcaaaaattttgcctcaaaactggtg<br>agctgaatttttgctgttaaagcatcgtgtagtgtttttcttagtccgttatgtaggtaggaatctgatgtaa<br>tggttgttggtattttgtcaccattcatttttatctggttgttctcaagttcggttacgagatccatttgtctatct<br>agttcaacttggaaaatcaacgtatcagtcgggcggcctcgcttatcaaccaccaatttcatattgct<br>gtaagtgtttaaatctttacttattggtttcaaaacccattggttaagccttttaaactcatggtagttattttc<br>aagcattaacatgaacttaaattcatcaaggctaatctctatatttgccttgtgagttttcttttgtgttagtt<br>cttttaataaccactcataaatcctcatagagtatttgttttcaaaagacttaacatgttccagattatattt<br>tatgaatttttttaactggaaaagataaggcaatatctcttcactaaaaactaattctaatttttcgcttga<br>gaacttggcatagtttgtccactggaagatcttaaagcctttaaccaaaggattcctgatttccacagtt<br>ctcgtcatcagctctctggttgctttagctaatacaccataagcattttccctactgatgttcatcatctga<br>gcgtattggttataagtgaacgataccgtccgttctttccttgtagggttttcaatcgtggggttgagtagt<br>gccacacagcataaaattagcttggtttcatgctccgttaagtcatagcgactaatcgctagttcatttg<br>ctttgaaaacaactaattcagacatacatctcaattggtctaggtgattttaatcactataccaattgag<br>atgggctagtcaatgataattactagtccttttcctttgagttgtgggtatctgtaaattctgctagaccttt<br>gctggaaaacttgtaaattctgctagaccctctgtaaattccgctagacctttgtgtgttttttttgtttatatt<br>caagtggttataatttatagaataaagaaagaataaaaaaagataaaaagaatagatcccagcc<br>ctgtgtataactcactactttagtcagttccgcagtattacaaaaggatgtcgcaaacgctgtttgctcc<br>tctacaaaacagacctGGCGCGCCCAGCTGTCTAGGGCGGCGGATTTGT<br>CCTACTCAGGAGAGCGTTCACCGACAAACAACAGATAAAACGAAA<br>GGCCCAGTCTTTCGACTGAGCCTTTCGTTTTATTTGATGCCTTTAAT<br>TAA |
| sTR060<br><br>[PrT7-triggerRNA] | taatacgactcactataggcataGCAGGGATAAACGAGATAGATAAGATAAGA<br>cccgctagcataacccttggggcctctaaacgggtcttgaggggttttttgctgaaaggaggaact<br>atatccggattggcgaatgggacgcgccctgtagcggcgcattaagcgcggcgggtgtggtggtt<br>acgcgcagcgtgaccgctacacttgccagcgccctagcgcccgctcctttcgctttcttcccttcctttc<br>tcgccacgttcgccggctttccccgtcaagctctaaatcgggggctccctttagggttccgatttagtg<br>ctttacggcacctcgaccccaaaaaacttgattagggtgatggttcacgtagtgggccatcgccctg<br>atagacggtttttcgccctttgacgttggagtccacgttctttaatagtggactcttgttccaaactggaa<br>caacactcaaccctatctcggtctattcttttgatttataagggattttgccgatttcggcctattggttaaa<br>aaatgagctgatttaacaaaaatttaacgcgaattttaacaaaatattaacgtttacaatttcaggtgg<br>cacttttcggggaaatgtgcgcggaacccctatttgtttatttttctaaatacattcaaatatgtatccgct<br>catgagacaataaccctgataaatgcttcaataatattgaaaaaggaagagtatgagtattcaaca<br>tttccgtgtcgcccttattcccttttttgcggcattttgccttcctgtttttgctcacccagaaacgctggtga<br>aagtaaaagatgctgaagatcagttgggtgcacgagtgggttacatcgaactggatctcaacagc |

| | |
|---|---|
| | ggtaagatccttgagagttttcgccccgaagaacgttttccaatgatgagcacttttaaagttctgctat<br>gtggcgcggtattatcccgtattgacgccgggcaagagcaactcggtcgccgcatacactattctca<br>gaatgacttggttgagtactcaccagtcacagaaaagcatcttacggatggcatgacagtaagag<br>aattatgcagtgctgccataaccatgagtgataacactgcggccaacttacttctgacaacgatcgg<br>aggaccgaaggagctaaccgcttttttgcacaacatgggggatcatgtaactcgccttgatcgttgg<br>gaaccggagctgaatgaagccataccaaacgacgagcgtgacaccacgatgcctgcagcaat<br>ggcaacaacgttgcgcaaactattaactggcgaactacttactctagcttcccggcaacaattaata<br>gactggatggaggcggataaagttgcaggaccacttctgcgctcggcccttccggctggctggttta<br>ttgctgataaatctggagccggtgagcgtgggtctcgcggtatcattgcagcactggggccagatg<br>gtaagccctcccgtatcgtagttatctacacgacggggagtcaggcaactatggatgaacgaaata<br>gacagatcgctgagataggtgcctcactgattaagcattggtaactgtcagaccaagtttactcatat<br>atactttagattgatttaaaacttcatttttaatttaaaaggatctaggtgaagatcctttttgataatctcat<br>gaccaaaatcccttaacgtgagttttcgttccactgagcgtcagaccccgtagaaaagatcaaagg<br>atcttcttgagatcctttttttctgcgcgtaatctgctgcttgcaaacaaaaaaaccaccgctaccagcg<br>gtggtttgtttgccggatcaagagctaccaactctttttccgaaggtaactggcttcagcagagcgca<br>gataccaaatactgtccttctagtgtagccgtagttaggccaccacttcaagaactctgtagcaccgc<br>ctacatacctcgctctgctaatcctgttaccagtggctgctgccagtggcgataagtcgtgtcttaccg<br>ggttggactcaagacgatagttaccggataaggcgcagcggtcgggctgaacggggggttcgtg<br>cacacagcccagcttggagcgaacgacctacaccgaactgagatacctacagcgtgagctatga<br>gaaagcgccacgcttcccgaagggagaaaggcggacaggtatccggtaagcggcagggtcgg<br>aacaggagagcgcacgagggagcttccagggggaaacgcctggtatctttatagtcctgtcgggt<br>ttcgccacctctgacttgagcgtcgatttttgtgatgctcgtcaggggggcggagcctatggaaaaac<br>gccagcaacgcggcctttttacggttcctggccttttgctggccttttgctcacatgttctttcctgcgttat<br>cccctgattctgtggataaccgtattaccgcctttgagtgagctgataccgctcgccgcagccgaac<br>gaccgagcgcagcgagtcagtgagcgaggaagcggaagagcgcctgatgcggtattttctcctta<br>cgcatctgtgcggtatttcacaccgcatatatggtgcactctcagtacaatctgctctgatgccgcata<br>gttaagccagtatacactccgctatcgctacgtgactgggtcatggctgcgccccgacacccgcca<br>acacccgctgacgcgccctgacgggcttgtctgctcccggcatccgcttacagacaagctgtgacc<br>gtctccgggagctgcatgtgtcagaggttttcaccgtcatcaccgaaacgcgcgaggcagctgcgg<br>taaagctcatcagcgtggtcgtgaagcgattcacagatgtctgcctgttcatccgcgtccagctcgttg<br>agtttctccagaagcgttaatgtctggcttctgataaagcgggccatgttaagggcggttttttcctgttt<br>ggtcactgatgcctccgtgtaagggggatttctgttcatgggggtaatgataccgatgaaacgagag<br>aggatgctcacgatacgggttactgatgatgaacatgcccggttactggaacgttgtgagggtaaa<br>caactggcggtatggatgcggcgggaccagagaaaaatcactcagggtcaatgccagcgcttcg<br>ttaatacagatgtaggtgttccacagggtagccagcagcatcctgcgatgcagatccggaacataa<br>tggtgcagggcgctgacttccgcgtttccagactttacgaaacacggaaaccgaagaccattcatg<br>ttgttgctcaggtcgcagacgttttgcagcagcagtcgcttcacgttcgctcgcgtatcggtgattcattc<br>tgctaaccagtaaggcaaccccgccagcctagccgggtcctcaacgacaggagcacgatcatg<br>cgcacccgtggggccgccatgccggcgataatggcctgcttctcgccgaaacgtttggtggcggg<br>accagtgacgaaggcttgagcgagggcgtgcaagattccgaataccgcaagcgacaggccgat<br>catcgtcgcgctccagcgaaagcggtcctcgccgaaaatgacccagagcgctgccggcacctgt<br>cctacgagttgcatgataaagaagacagtcataagtgcggcgacgatagtcatgccccgcgccc<br>accggaaggagctgactgggttgaaggctctcaagggcatcggtcgagatcccggtgcctaatga<br>gtgagctaacttacattaattgcgttgcgctcactgcccgctttccagtcgggaaacctgtcgtgccag<br>ctgcattaatgaatcggccaacgcgcggggagaggcggtttgcgtattgggcgccagggtggttttt<br>cttttcaccagtgagacgggcaacagctgattgcccttcaccgcctggccctgagagagttgcagc<br>aagcggtccacgctggtttgccccagcaggcgaaaatcctgtttgatggtggttaacggcgggatat<br>aacatgagctgtcttcggtatcgtcgtatcccactaccgagatatccgcaccaacgcgcagcccgg<br>actcggtaatggcgcgcattgcgcccagcgccatctgatcgttggcaaccagcatcgcagtggga<br>acgatgccctcattcagcatttgcatggtttgttgaaaaccggacatggcactccagtcgccttcccgt<br>tccgctatcggctgaatttgattgcgagtgagatatttatgccagccagccagacgcagacgcgcc<br>gagacagaacttaatgggcccgctaacagcgcgatttgctggtgacccaatgcgaccagatgctc<br>cacgcccagtcgcgtaccgtcttcatgggagaaaataatactgttgatgggtgtctggtcagagaca<br>tcaagaaataacgccggaacattagtgcaggcagcttccacagcaatggcatcctggtcatccag |

| | |
|---|---|
| | cggatagttaatgatcagcccactgacgcgttgcgcgagaagattgtgcaccgccgctttacaggc<br>ttcgacgccgcttcgttctaccatcgacaccaccacgctggcacccagttgatcggcgcgagattta<br>atcgccgcgacaatttgcgacggcgcgtgcagggccagactggaggtggcaacgccaatcagc<br>aacgactgtttgcccgccagttgttgtgccacgcggttgggaatgtaattcagctccgccatcgccgc<br>ttccactttttcccgcgttttcgcagaaacgtggctggcctggttcaccacgcgggaaacggtctgata<br>agagacaccggcatactctgcgacatcgtataacgttactggtttcacattcaccaccctgaattgac<br>tctcttccgggcgctatcatgccataccgcgaaaggttttgcgccattcgatggtgtccgggatctcga<br>cgctctcccttatgcgactcctgcattaggaagcagcccagtagtaggttgaggccgttgagcaccg<br>ccgccgcaaggaatggtgcatgcaaggagatggcgcccaacagtcccccggccacggggcct<br>gccaccatacccacgccgaaacaagcgctcatgagcccgaagtggcgagcccgatcttccccat<br>cggtgatgtcggcgatataggcgccagcaaccgcacctgtggcgccggtgatgccggccacgat<br>gcgtccggcgtagaggatcgagatctcgatcccgcgaaat |
| *C. jejuni* HipO Gblocks<br><br>[Pr1-HipO CDS] | ctcggatacccttactctgttgaaaacgaatagataggtttgagctaacaccgtgcgtgttgacaatttt<br>acctctggcggtgataatggttgcaAATAATTTTGTTTAACTTTAAGAAGGAGAT<br>ATACCATGAACCTGATCCCGGAAATCCTGGACCTGCAGGGTGAATT<br>CGAAAAAATCCGTCACCAGATCCACGAAAACCCGGAACTGGGTTT<br>CGACGAACTGTGCACCGCTAAACTGGTTGCTCAGAAACTGAAAGA<br>ATTCGGTTACGAAGTTTACGAAGAAATCGGTAAAACCGGTGTTGTT<br>GGTGTTCTGAAAAAAGGTAACTCTGACAAAAAAATCGGTCTGCGTG<br>CTGACATGGACGCTCTGCCGCTGCAGGAATGCACCAACCTGCCGT<br>ACAAATCTAAAAAAGAAAACGTTATGCACGCTTGCGGTCACGACGG<br>TCACACCACCTCTCTGCTGCTGGCTGCTAAATACCTGGCTTCTCAG<br>AACTTCAACGGTGCTCTGAACCTGTACTTCCAGCCGGCTGAAGAA<br>GGTCTGGGTGGTGCTAAAGCTATGATCGAAGACGGTCTGTTCGAA<br>AAATTCGACTCTGACTACGTTTTCGGTTGGCACAACATGCCGTTCG<br>GTTCTGACAAAAAATTCTACCTGAAAAAAGGTGCTATGATGGCTTCT<br>TCTGACTCTTACTCTATCGAAGTTATCGGTCGTGGTGGTCACGGTT<br>CTGCTCCGGAAAAAGCTAAAGACCCGATCTACGCTGCTTCTCTGCT<br>GATCGTTGCTCTGCAGTCTATCGTTTCTCGTAACGTTGACCCGCAG<br>AACTCTGCTGTTGTTTCTATCGGTGCTTTCAACGCTGGTCACGCTT<br>TCAACATCATCCCGGACATCGCTACCATCAAAATGTCTGTTCGTGC<br>TCTGGACAACGAAACCCGTAAACTGACCGAAGAAAAAATCTACAAA<br>ATCTGCAAAGGTATCGCTCAGGCTAACGACATCGAAATCAAAATCA<br>ACAAAAACGTTGTTGCTCCGGTTACCATGAACAACGACGAAGCTGT<br>TGACTTCGCTTCTGAAGTTGCTAAAGAACTGTTCGGTGAAAAAAAC<br>TGCGAATTCAACCACCGTCCGCTGATGGCTTCTGAAGACTTCGGTT<br>TCTTCTGCGAAATGAAAAAATGCGCTTACGCTTTCCTGGAAAACGA<br>AAACGACATCTACCTGCACAACTCTTCTTACGTTTTCAACGACAAAC<br>TGCTGGCTCGTGCTGCTTCTTACTACGCTAAACTGGCTCTGAAATA<br>CCTGAAATAATAAattattgaccacttccgagtagaatcgtgcttcagtaaga |
| *H. ailurogastricus* HipO Gblocks<br><br>[Pr1-HipO CDS] | ctcggatacccttactctgttgaaaacgaatagataggtttgagctaacaccgtgcgtgttgacaatttt<br>acctctggcggtgataatggttgcaAATAATTTTGTTTAACTTTAAGAAGGAGAT<br>ATACCATGCCGCTGATCCCGGAAATCGTTGCTATGCAGGAAGAATT<br>CCAGGCTATCCGTCAGCAGATCCACCAGGACCCGGAACTGGGTTT<br>CGAAGAAGTTCTGACCTCTGGTCTGGTTGCTGACAAACTGAAAGAA<br>TTCGGTTACGAAGTTCACACCGGTGTTGGTAAAACCGGTGTTGTTG<br>GTGTTCTGAAAAAAGGTAACCTGGCTAAAAAAATCGGTCTGCGTGC<br>TGACATGGACGCTCTGCCGATGCCGGAACACAACGACCTGCCGTA<br>CAAATCTCAGATCCCGAACCGTATGCACGCTTGCGGTCACGACGG<br>TCACTCTGCTTCTCTGCTGCTGGCTGCTAAATACCTGGCTTCTCAG<br>GAATTCAACGGTATCCTGAACCTGTACCTGCAGCCGGCTGAAGAA<br>GGTCTGGGTGGTGCTAAAGCTATGCTGGAAGACGGTCTGCTGGAA |

| | |
|---|---|
| | CGTTTCGACTCTGACATGATCTTCGGTTGGCACAACATCCCGCTGG<br>GTACCGACAAAAAAATCTACCTGGAATCTGGTGCTGTTATGGCTTC<br>TGCTGACTCTTACACCATCGAAATCAAAGGTCAGGGTGGTCACGGT<br>TCTGCTCCGGAAAAATGCAAAGACCCGGTTCTGGCTGCTTCTCTGC<br>TGGTTGTTGCTCTGCAGTCTATCGTTTCTCGTAACATCGACCCGCA<br>GCACTCTGCTGTTGTTTCTGTTGGTGCTTTCAACGCTGGTAACACC<br>TTCAACATCATCCCGGACCGTGCTACCCTGAAACTGTCTGTTCGTG<br>CTCTGGACGCTGAATCTCAGGAAATCGTTGCTGAACACATCTCTAA<br>AATCGCTAAAGGTATCGCTCTGGCTCACGGTGTTGAAATCGAAATC<br>ACCAAACAGGCTGCTGCTACCATCATGTTCAACGACCCGAAAGCTA<br>CCGCTTTCGCTCAGGAAGTTGCTCTGGAAGTTTTCGGTAAAGAAGC<br>TTGCTGCTTCGAATACCCGGCTGCTATGGGTTCTGAAGACTTCGGT<br>TACTTCGCTCAGCTGCGTCCGTGCGCTTACGCTTTCCTGGAAAACG<br>AAAACACCCACTACCTGCACACCTCTTCTTACGTTTTCAACGACGC<br>TCTGCTGGCTCGTGCTGCTTCTTACTACGCTCGTCTGGTTCTGAAC<br>TACCTGAAATAATAAattattgaccacttccgagtagaatcgtgcttcagtaaga |
| *S. enterica* HipO Gblocks<br><br>[Pr1-HipO CDS] | ctcggatacccttactctgttgaaaacgaatagataggtttgagctaacaccgtgcgtgttgacaatttt<br>acctctggcggtgataatggttgcaAATAATTTTGTTTAACTTTAAGAAGGAGAT<br>ATACCATGAAACTGATCAACGAAATCGTTAAAACCCAGAAAGAATT<br>CGCTTCTGTTCGTCAGAAAATCCACAAAAACCCGGAACTGGGTTTC<br>CAGGAAGTTGCTACCGCTAAACTGGTTGCTGGTCTGCTGGGTGAA<br>TACGGTTACCAGGTTCACGAAAAAGTTGGTCGTACCGGTGTTGTTG<br>CTGTTCTGAAAAAAGGTAACGGTAACAAAAAAATCGGTCTGCGTGC<br>TGACATGGACGCTCTGCCGATGCAGGAACTGGCTGAAGTTCCGTA<br>CAAATCTGTTATCCCGGGTGTTATGCACGCTTGCGGTCACGACGG<br>TCACACCGCTTCTCTGCTGATGGCTGCTAAATACCTGTCTCAGTGC<br>GACTTCAACGGTCAGCTGAACCTGATCTTCCAGCCGGCTGAAGAA<br>GGTTCTGGTGGTGCTCTGTCTATGATCAACGACGGTCTGTTCGAAC<br>GTTTCGACTGCGACTACATCTTCTCTTGGCACAACCTGCCGTGCAA<br>AAACCAGGACAAACAGATCTTCCTGAAAAAAGGTGTTTTCCTGTCT<br>TCTTCTGACCGTTTCAAAATCAAAATCGCTGGTTCTGGTGGTCACG<br>CTTCTGCTCCGCAGAACTCTAAAGACCCGACCCTGGCTGCTTGCC<br>ACCTGATCCTGGCTCTGCAGTCTATCGTTTCTCGTAACACCGACCC<br>GCAGCAGTCTGTTGTTATCTCTGTTGGTTCTATCATCGCTGGTAAC<br>GACGAATCTTACAACATCATCCCGGAACAGGTTGAAATCCTGCTGT<br>CTGTTCGTACCCTGAACAAAAACGTTCGTAAACAGACCATCAAACG<br>TATCAACGAAATCATCGAACACTGCTCTAACCTGTTCGGTCTGACC<br>TCTGAAGTTGAATACTACGACAAAGCTGACGTTACCTACAACGACG<br>AAGAAGCTACCTCTCTGGCTTGGAAAGTTGCTGGTGAAATCTTCGG<br>TAACGAATGCTGCGCTTTCGAACACTCTCCGGGTATGGCTTCTGAC<br>GACCTGTCTTACATGCTGTCTGCTCGTAAAGGTTGCTACGCTTACA<br>TCAACAACGGTGACACCGCTTACGTTCACAACGGTCACTACGTTTT<br>CAACGACGACCTGCTGTCTATCGCTGCTACCTACTTCGCTAAAATC<br>ACCCTGGAATACCTGCAGTAATAAattattgaccacttccgagtagaatcgtgcttca<br>gtaaga |
| *H. felis* HipO Gblocks<br><br>[Pr1-HipO CDS] | ctcggatacccttactctgttgaaaacgaatagataggtttgagctaacaccgtgcgtgttgacaatttt<br>acctctggcggtgataatggttgcaAATAATTTTGTTTAACTTTAAGAAGGAGAT<br>ATACCATGAACCTGATCCCGGAAATCGTTGCTATGCAGGAAGAATT<br>CATCGCTATCCGTCACCAGATCCACCGTCACCCGGAACTGGGTTT<br>CAAAGAAGTTCAGACCTCTCAGCTGGTTGCTGACAAACTGCGTGAA<br>TTCGGTTACGAAGTTCACACCGGTGTTGGTAAAACCGGTGTTGTTG<br>GTGTTCTGAAAAAAGGTGACTCTGCTAAAAAAATCGGTCTGCGTGC |

| | |
|---|---|
| | TGACATGGACGCTCTGCCGATCCAGGAAGACTCTGGTCTGGACTA<br>CCAGTCTCAGACCCCGCAGCGTATGCACGCTTGCGGTCACGACGG<br>TCACTCTGCTTCTCTGCTGCTGGCTGCTAAATACCTGGCTACCCAG<br>GACTTCAAAGGTACCCTGCACCTGTACTTCCAGCCGGCTGAAGAA<br>AACCTGGGTGGTGCTAAAGCTATGATCGAAGAAGGTCTGCTGGAA<br>AAATTCGACTCTGACCTGATCTTCGGTTGGCACAACATGCCGCTGG<br>GTTCTGACAAAAAATTCTACCTGAAATCTGGTGCTATGATGGCTTCT<br>TCTGACGCTTACACCCTGGAAATCAAAGCTCAGGGTGGTCACGCTT<br>CTGCTCCGGAAAAAACCAAAGACCCGATCCTGGTTGCTTCTCTGCT<br>GGTTCTGGCTCTGCAGGGTATCATCTCTCGTAACGTTGACCCGCA<br>GAACTCTGCTGTTGTTTCTGTTGGTGCTCTGAACGCTGGTTCTGCT<br>CACAACATCATCCCGGACCGTGCTGTTCTGCTGGTTTCTGTTCGTG<br>CTCTGGACTCTCTGACCCGTGAACTGGTTGCTAAACGTATCCAGGA<br>AATCTGCCAGGGTGTTGCTCTGGCTCAGGGTGTTGAAATCAACATC<br>ACCCACGAATTCGCTACCCCGATCACCAACAACCACTCTGAAGCTA<br>CCGCTCTGGCTCAGGAAGTTGCTCTGGACATCTTCGGTGCTCAGG<br>ACTGCTGCTTCGACCACAAACCGGCTATGGGTTCTGAAGACTTCG<br>GTTACTTCTGCGAACGTCGTAAATGCGCTTACGCTTTCTTCGAAAA<br>CGAAACCACCCACTACATCCACACCTCTAACTACGTTTTCAACGAC<br>GCTCTGCTGGCTCGTGCTGCTTCTTACTACGCTGGTCTGGTTCTGA<br>AATACCTGGGTTAATAAattattgaccacttccgagtagaatcgtgcttcagtaaga |
| *C. coli* HipO Gblocks<br><br>[Pr1-HipO CDS] | ctcggatacccttactctgttgaaaacgaatagataggtttgagctaacaccgtgcgtgttgacaatttt<br>acctctggcggtgataatggttgcaAATAATTTTGTTTAACTTTAAGAAGGAGAT<br>ATACCATGAACCTGATCCCGGAAATCCTGGACCTGCAGGGTGAATT<br>CGAAAAAATCCGTCACCAGATCCACGAAAACCCGGAACTGGGTTT<br>CGACGAACTGTGCACCGCTAAACTGGTTGTTCAGAAACTGAAAGAA<br>TTCGGTTACGAAGTTTACGAAGAAATCGGTAAAACCGGTGTTGTTG<br>GTGTTCTGAAAAAAGGTAACTCTGACAAAAAAATCGGTCTGCGTGC<br>TGACATGGACGCTCTGCCGCTGCAGGAATACACCAACCTGCCGTA<br>CAAATCTAAAAAAGAAAACGTTATGCACGCTTGCGGTCACGACGGT<br>CACACCACCTCTCTGCTGCTGGCTGCTAAATACCTGGCTTCTCAGA<br>ACTTCAACGGTACCCTGAACATCTACTTCCAGCCGGCTGAAGAAG<br>GTCTGGGTGGTGCTAAAGCTATGATCGAAGACGGTCTGTTCGAAA<br>AATTCGACTCTGACTACGTTTTCGGTTGGCACAACATGCCGTTCGG<br>TTCTGACAAAAAATTCTACCTGAAAAAAGGTGCTATGATGGCTTCTT<br>CTGACTCTTACTCTATCGAAGTTATCGGTCGTGGTGGTCACGGTTC<br>TGCTCCGGAAAAAGCTAAAGACCCGATCTACGCTGCTTCTCTGCTG<br>GTTGTTGCTCTGCAGTCTATCGTTTCTCGTAACGTTGACCCGCAGA<br>ACTCTGCTGTTGTTTCTATCGGTGCTTTCAACGCTGGTTACGCTTTC<br>AACATCATCCCGGACATCGCTATGATCAAAATGTCTGTTCGTGCTC<br>TGGACAACGAAACCCGTAAACTGACCGAAGAAAAAATCTACAAAAT<br>CTGCAAAGGTATCGCTCAGGCTAACGACATCGGTATCAAAATCAAC<br>AAAAACGTTGTTGCTCCGGTTACCATGAACAACGACGAAGCTGTTG<br>ACTTCGCTTCTGAAGTTGCTAAAGAACTGTTCGGTGAAAAAAACTG<br>CGAATTCAACCACCGTCCGCTGATGGCTTCTGAAGACTTCGGTTTC<br>TTCTGCGAAATGAAAAAATGCGCTTACGCTTTCCTGGAAAACGAAA<br>ACGACATCTACCTGCACAACTCTTCTTACGTTTTCAACGACAAACTG<br>CTGGCTCGTGCTGCTTCTTACTACGCTAAACTGGCTCTGAAATACC<br>TGAAATAATAAattattgaccacttccgagtagaatcgtgcttcagtaaga |

### Supplementary Table 6

Descriptions of which data comes from which CFS conditions are presented below. The experimental conditions can be broadly grouped into those used at the Paris lab [green] and the Montpellier lab [blue].

| | | Figures | | | | | | | | | | | | | | | | | | | | | | |
|---|---|---|---|---|---|---|---|---|---|---|---|---|---|---|---|---|---|---|---|---|---|---|---|---|
| **Condition** | **Variable** | 1 D | 2 B | 2 C | 2 D | 3 A | 3 B | 4 A | 4 B | 5 C | 6 D | S 3 | S 4 A | S 4 B | S 5 B | S 5 C | S 5 D | S 6 | S 7 A | S 7 B | S 8 B | S 8 C | S 8 D | S 9 |
| Lysis method | Sonication | green | | | | green | green | | | | | green | | | green | green | green | | | | green | green | green | |
| | French Press | | blue | blue | blue | | | blue | blue | blue | blue | | blue | blue | | | | blue | blue | blue | | | | blue |
| DNA concentration | 1 nM | | | | | green | green | | | | | | | | green | green | | | | | green | green | | |
| | 5 nM | green | | | | | | | | | | green | | | | | green | | | | | | green | |
| | 4 nM | | blue | blue | | | | | | | | | | | | | | | | | | | | |
| | 10 nM | | | | blue | | | blue | blue | blue | blue | | | | | | | blue | blue | blue | | | | blue |
| Reaction volume | 10.5 µL | green | | | | green | green | | | | | green | | | green | green | green | | | | green | green | green | |
| | 20 µL | | blue | blue | blue | | | blue | blue | blue | blue | | blue | blue | | | | blue | blue | blue | | | | blue |
| Temperature | 37C | | blue | blue | blue | | | blue | blue | blue | blue | | blue | blue | | | | blue | blue | blue | | | | blue |
| | 30C | green | | | | green | green | | | | | green | | | green | green | green | | | | green | green | green | |
| Genetic background (see Supplementary Table 1) | BL21 | | blue | blue | blue | | | blue | | | | | | | | | | | blue | blue | | | | blue |
| | BL21 Δ*recB* | blue | | | | | | | | | | blue | | | | | | | | | | | | |
| | BL21 Δ*recBCD* | blue | blue | blue | blue | | | blue | blue | blue | blue | blue | blue | blue | | | | blue | | | | | | |
| | BL21 Rosetta2 | green | | | | green | green | | | | | green | | | green | green | green | | | | | | green | |
| | BL21 Rosetta2 Δ*recB* | | | | | green | green | | | | | | | | green | green | green | | | | | | green | |
| | BL21 Rosetta2 Δ*recBCD* | | | | | green | green | | | | | | | | green | green | green | | | | | | green | |
| | BL21 Rosetta2 GamS | | | | | | | | | | | | | | | | | | | | green | green | green | |
| | BL21 Rosetta2 Δ*recB* GamS | | | | | | | | | | | | | | | | | | | | green | green | green | |
| | BL21 Rosetta2 Δ*recBCD* GamS | | | | | | | | | | | | | | | | | | | | green | green | green | |
| Construct | sfGFP | | blue | blue | blue | | | blue | blue | blue | blue | | blue | blue | | | | blue | blue | blue | | | | blue |
| | deGFP | green | | | | green | green | | | | | green | | | green | green | green | | | | green | green | green | |

| | | 1D | 2B | 2C | 2D | 3A | 3B | 4A | 4B | 5C | 6D | S3 | S4A | S4B | S5B | S5C | S5D | S6 | S7A | S7B | S8B | S8C | S8D | S9 |
|---|---|---|---|---|---|---|---|---|---|---|---|---|---|---|---|---|---|---|---|---|---|---|---|---|
| Flank length | 0 bp | | | | | | | | | | | | | | | | | | | | | | | |
| | 40 bp | | | | | | | | | | | | | | | | | | | | | | | |
| | 339 bp upstream and 212 bp downstream | | | | | | | | | | | | | | | | | | | | | | | |
| DNA Purification | Yes | | | | | | | | | | | | | | | | | | | | | | | |
| | No | | | | | | | | | | | | | | | | | | | | | | | |
| 384-well plate | Corning 384-well low volume black with clear bottom (#3540) | | | | | | | | | | | | | | | | | | | | | | | |
| | Nunc 384-well flat bottom (#142761) | | | | | | | | | | | | | | | | | | | | | | | |
| Fluorescence Plate Reader | Biotek Synergy H1M, | | | | | | | | | | | | | | | | | | | | | | | |
| | Biotek Cytation 3 | | | | | | | | | | | | | | | | | | | | | | | |
| | Biotek Synergy HTX plate reader | | | | | | | | | | | | | | | | | | | | | | | |

**Supplementary Table 7**
Primers used to amplify the linear DNA fragments are listed below:

| Fragment | Template | Forward primer^ | Reverse Primer^ |
|---|---|---|---|
| sfGFP | CMP | p33 | p34 |
| sfGFP-2MethoxyEtoxy | CMP | SP0_3 2-MethoxyEtoxy | SPN_3 2-MethoxyEtoxy |
| sfGFP-Phosphorothioate | CMP | SP0_1 Phosphorothioate | SPN_1 Phosphorothioate |
| sfGFP-FluoroC | CMP | SP0_5 FluoroC | SPN_5 FluoroC |
| sfGFP-Phosphorylation | CMP | SP0_2 Phosphorylation | SPN_2 Phosphorylation |
| deGFP | P70a-deGFP | p9 | p10 |
| P1-sw_Original RBS sTR056 | sTR056 | toehold_P1_add_FW | SBa_000587_RV |
| sw_J61100 | sTR056 | J61100_FW | SBa_000587_RV |
| sw_J61101 | sTR056 | J61101_FW | SBa_000587_RV |
| sw_J61102 | sTR056 | J61105_FW | SBa_000587_RV |
| sw_J61103 | sTR056 | J61106_FW | SBa_000587_RV |
| sw_J61104 | sTR056 | J61102_FW | SBa_000587_RV |
| sw_J61105 | sTR056 | J61103_FW | SBa_000587_RV |
| sw_J61106 | sTR056 | J61104_FW | SBa_000587_RV |
| sw_J61107 | sTR056 | J61109_FW | SBa_000587_RV |
| sw_J61108 | sTR056 | J61107_FW | SBa_000587_RV |
| sw_J61109 | sTR056 | J61108_FW | SBa_000587_RV |
| sw_J61110 | sTR056 | J61110_FW | SBa_000587_RV |
| P1-sw_J61100 | sw_J61100 | toehold_P1_add_FW | SBa_000587_RV |
| P1-sw_J61101 | sw_J61101 | toehold_P1_add_FW | SBa_000587_RV |
| P1-sw_J61102 | sw_J61102 | toehold_P1_add_FW | SBa_000587_RV |
| P1-sw_J61103 | sw_J61103 | toehold_P1_add_FW | SBa_000587_RV |
| P1-sw_J61104 | sw_J61104 | toehold_P1_add_FW | SBa_000587_RV |
| P1-sw_J61105 | sw_J61105 | toehold_P1_add_FW | SBa_000587_RV |
| P1-sw_J61106 | sw_J61106 | toehold_P1_add_FW | SBa_000587_RV |

| | | | |
|---|---|---|---|
| P1-sw_J61107 | sw_J61107 | toehold_P1_add_FW | SBa_000587_RV |
| P1-sw_J61108 | sw_J61108 | toehold_P1_add_FW | SBa_000587_RV |
| P1-sw_J61109 | sw_J61109 | toehold_P1_add_FW | SBa_000587_RV |
| P1-sw_J61110 | sw_J61110 | toehold_P1_add_FW | SBa_000587_RV |
| Cj_HipO | *C. jejuni* HipO Gblock | p33 | p34 |
| Ha_HipO | *H. ailurogastricus* HipO Gblock | p33 | p34 |
| Se_HipO | *S. enterica* HipO Gblock | p33 | p34 |
| Hf_HipO | *H. felis* HipO Gblock | p33 | p34 |
| Cc_HipO | *C. coli* HipO Gblock | p33 | p34 |

^Primer sequences are listed in Supplementary Table 3.
