## Supplementary Figures file for "Differentially optimized cell-free buffer enables robust expression from unprotected linear DNA in exonuclease-deficient extracts"

#### Fig S1: Colony PCR confirming ΔrecB and ΔrecBCD deletions

A

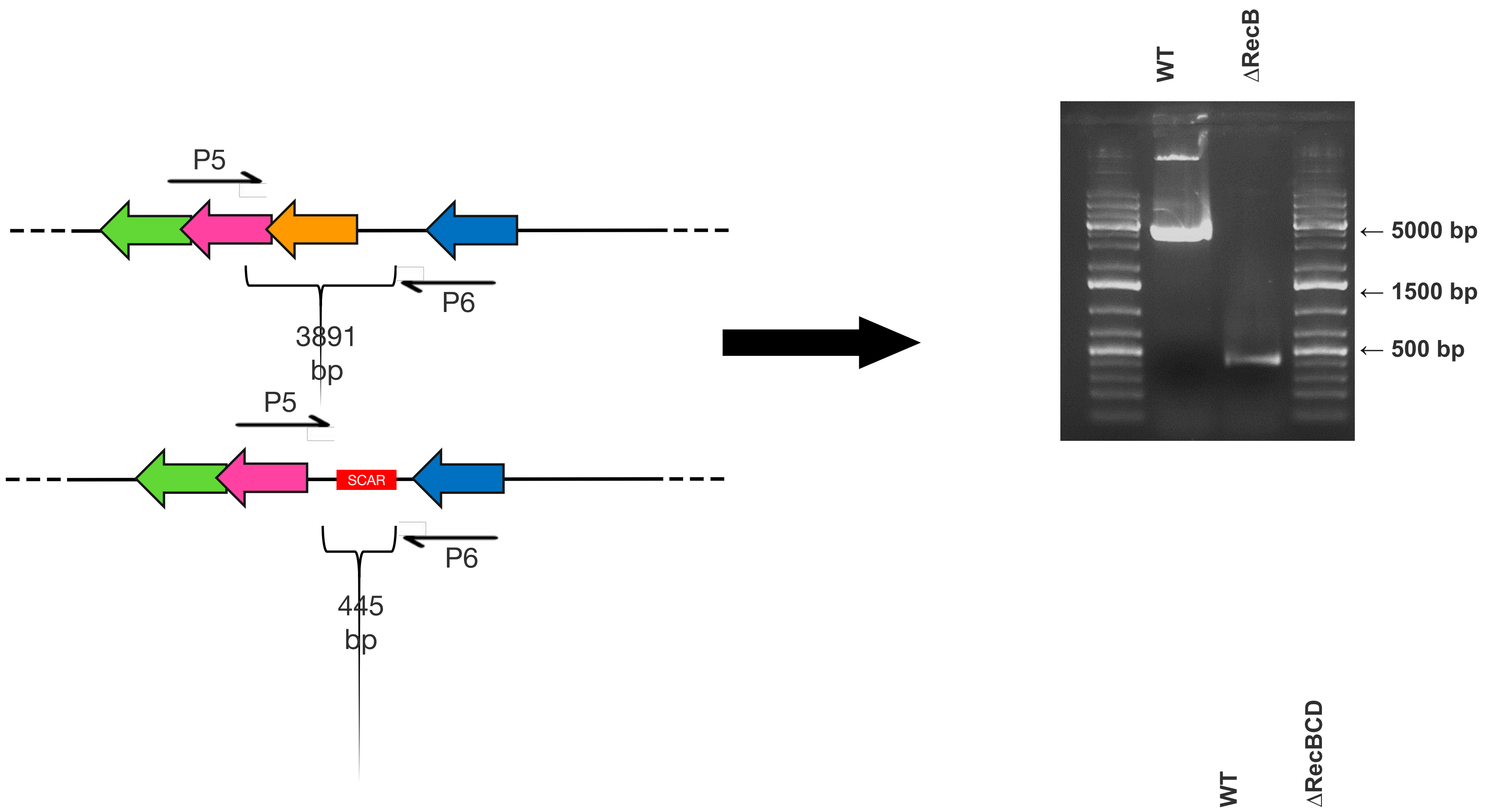


B

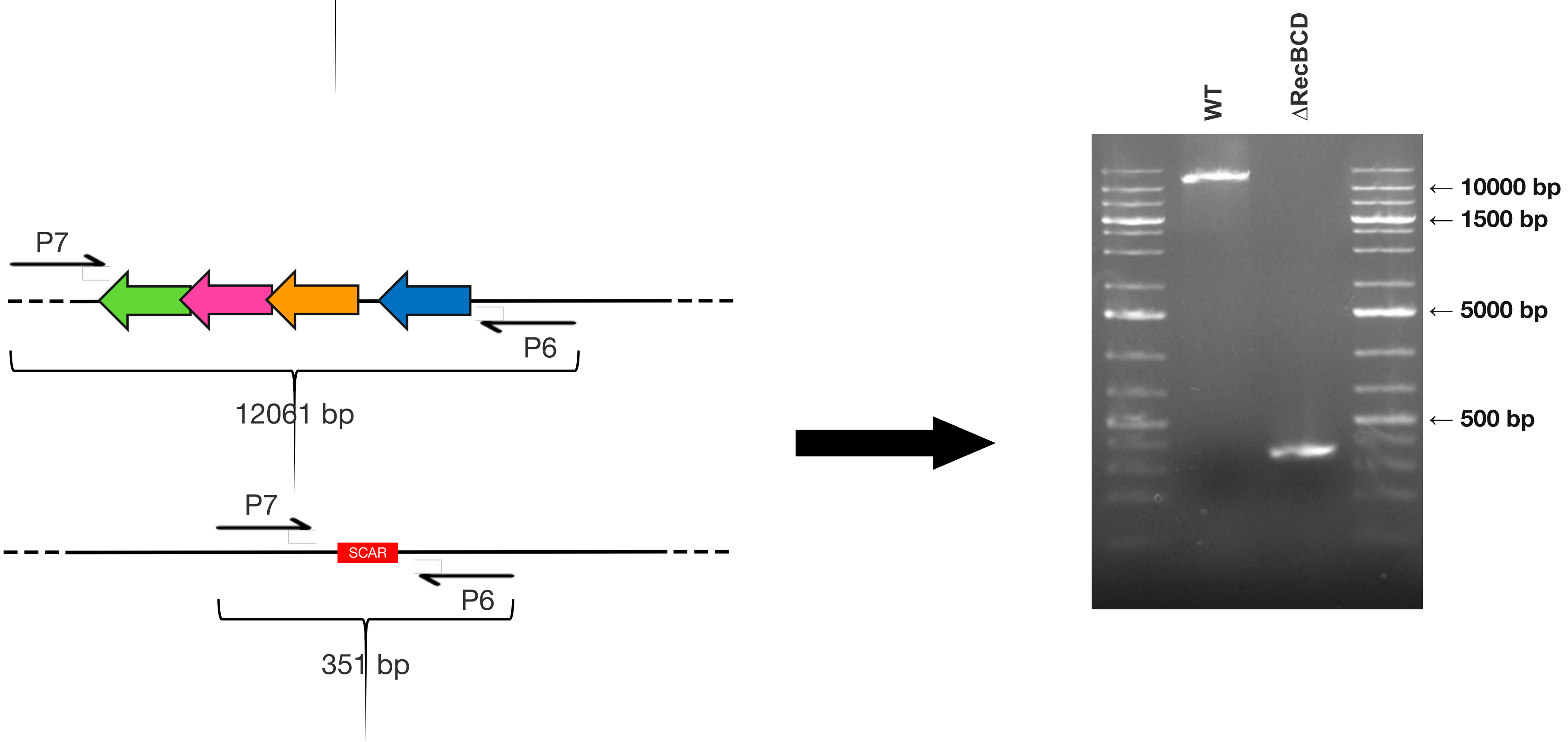


**Fig S1: Colony PCR confirming ΔrecB and ΔrecBCD deletions.** The genes deletion were confirmed by colony PCR after curing the residual Kan$^R$ cassette from homologous recombination. (A) *recB* gene was deleted, generating a scar of 445 bp that could be verified with primers P5 and P6 (left), and analyzed on a 1% agarose gel (right). (B) *recBCD* operon was deleted, generating a scar of 351 bp that could be verified with primers P7 and P8 (left), and analyzed on a 1% agarose gel (right).

### Fig S2: Specific growth rates of strains

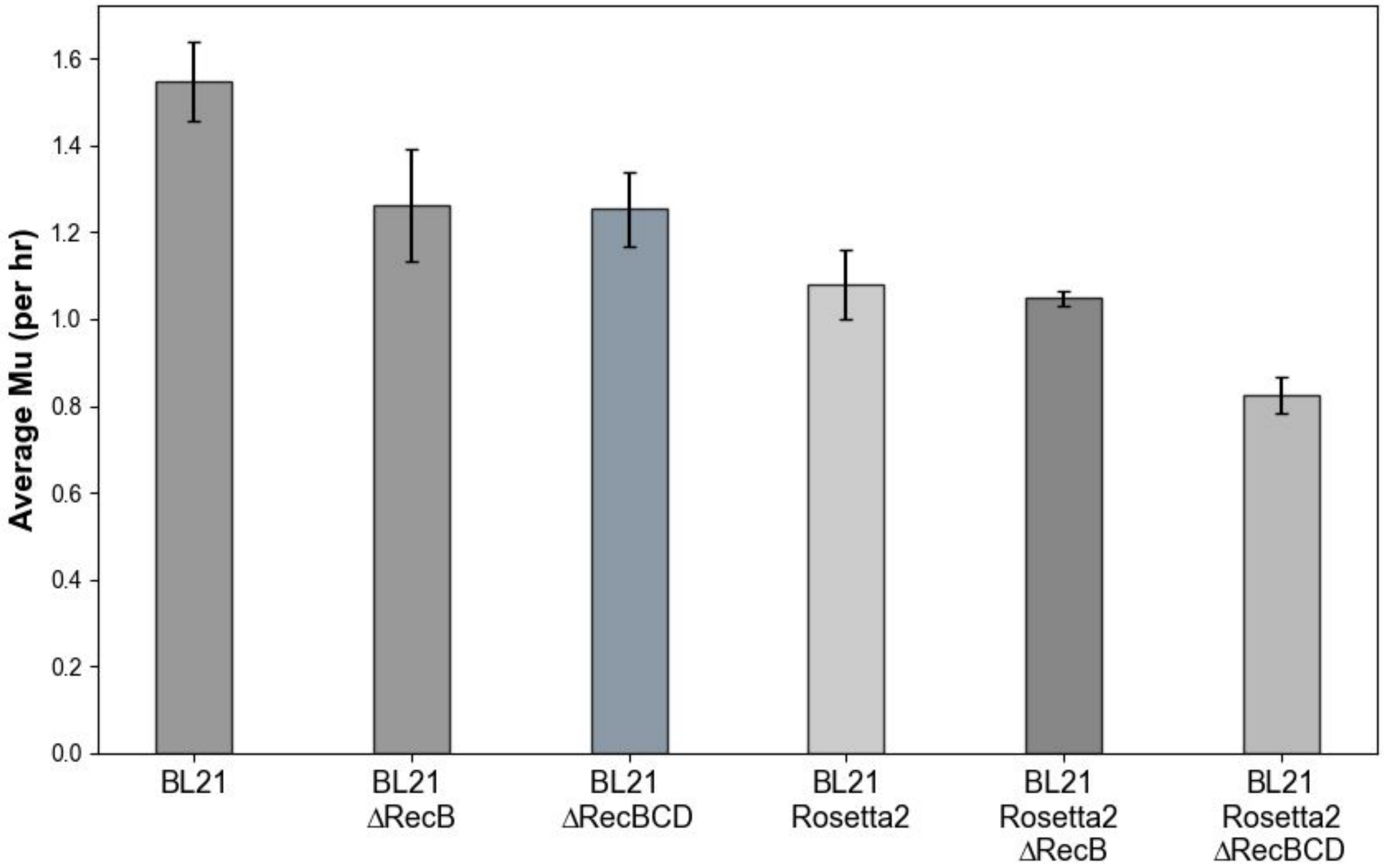


**Fig S2: Specific growth rates of strains.** All parental (BL21 and BL21 Rosetta2) and modified strains (BL21 *ΔrecB*, BL21 Δ*recBCD*, BL21 Rosetta2 Δ*recB*, and BL21 Rosetta2 Δ*recBCD*) were grown in 2YTP medium used for expanding cells for lysis. Specific growth rate of each strain was calculated in the log phase of growth from 4 colonies picked for inoculation. Compared to the parentals, strains did not show a substantial decrease in growth rate upon genomic deletions. Genotypes of the strains are listed in Supplementary Table 1. Data shown are mean±SD for 4 biological replicates done on the same day.

### Fig S3: 8h Kinetic curves for Fig 1D data

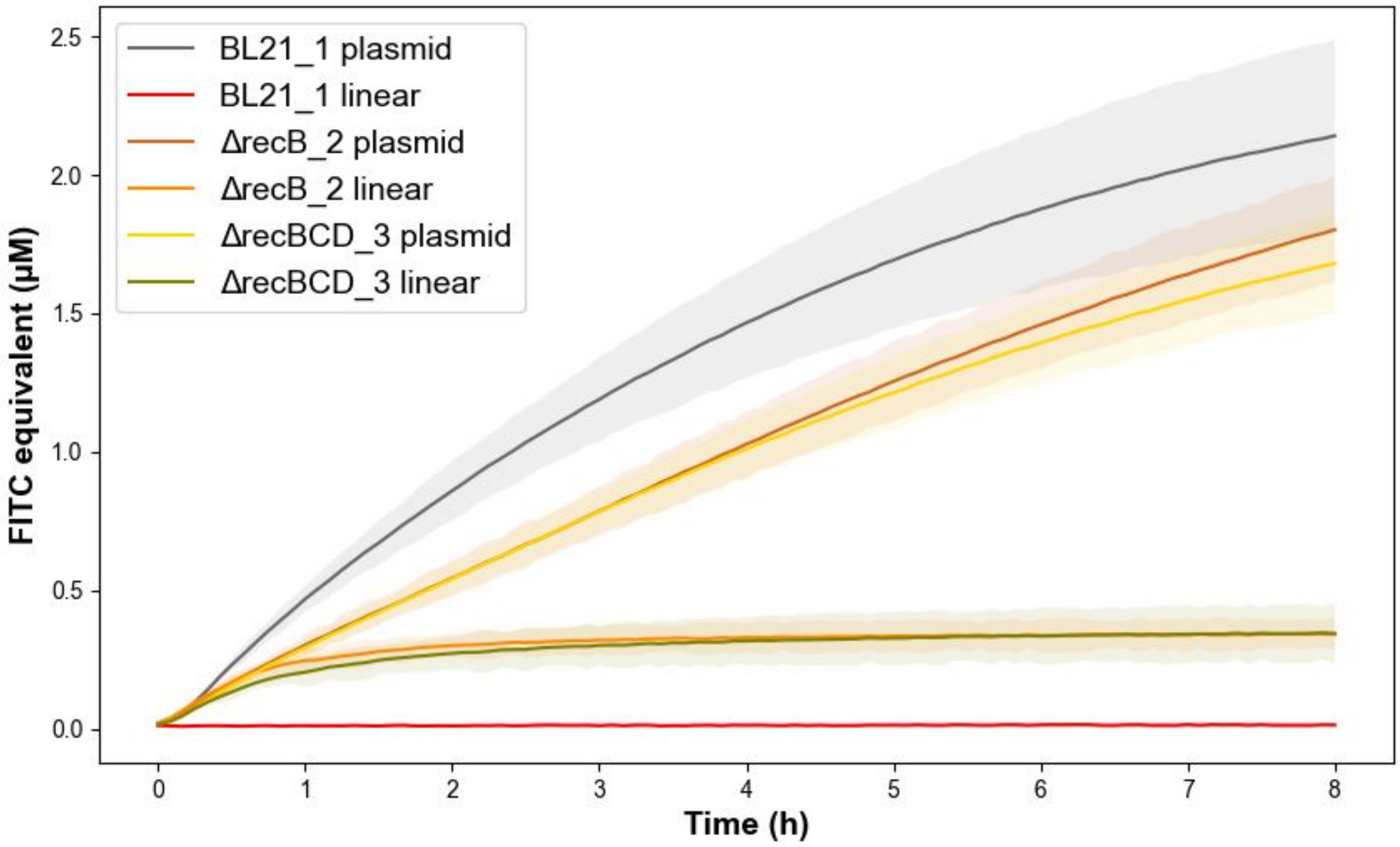


**Fig S3: 8h Kinetic curves for Fig 1D data.** Kinetic curves data from Figure 1D. Data shown are the mean±SD for 3 replicates done on the same day.

**Fig S4: Optimal linear and plasmid DNA concentration in Δ*recBCD* extracts.**

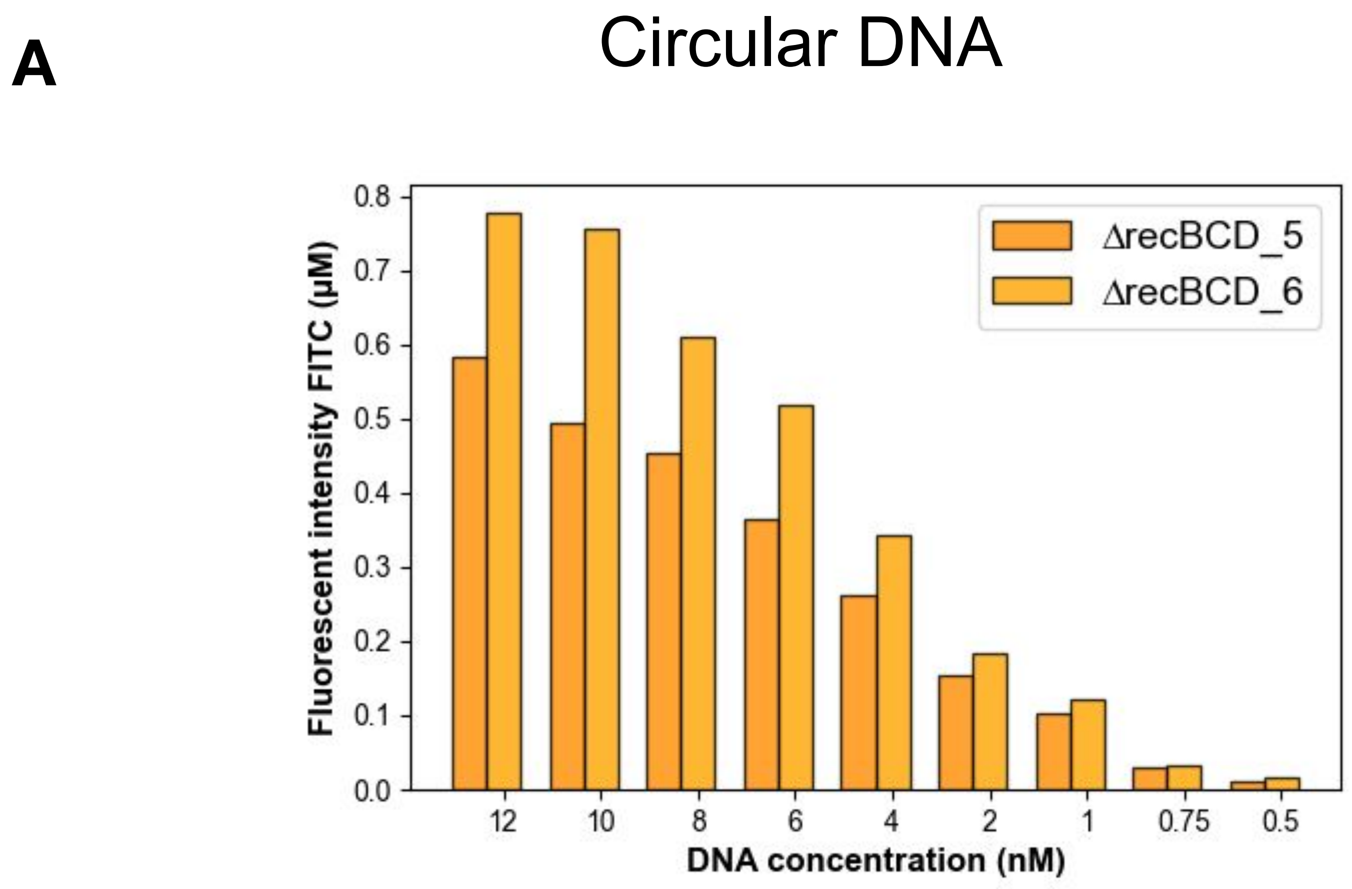


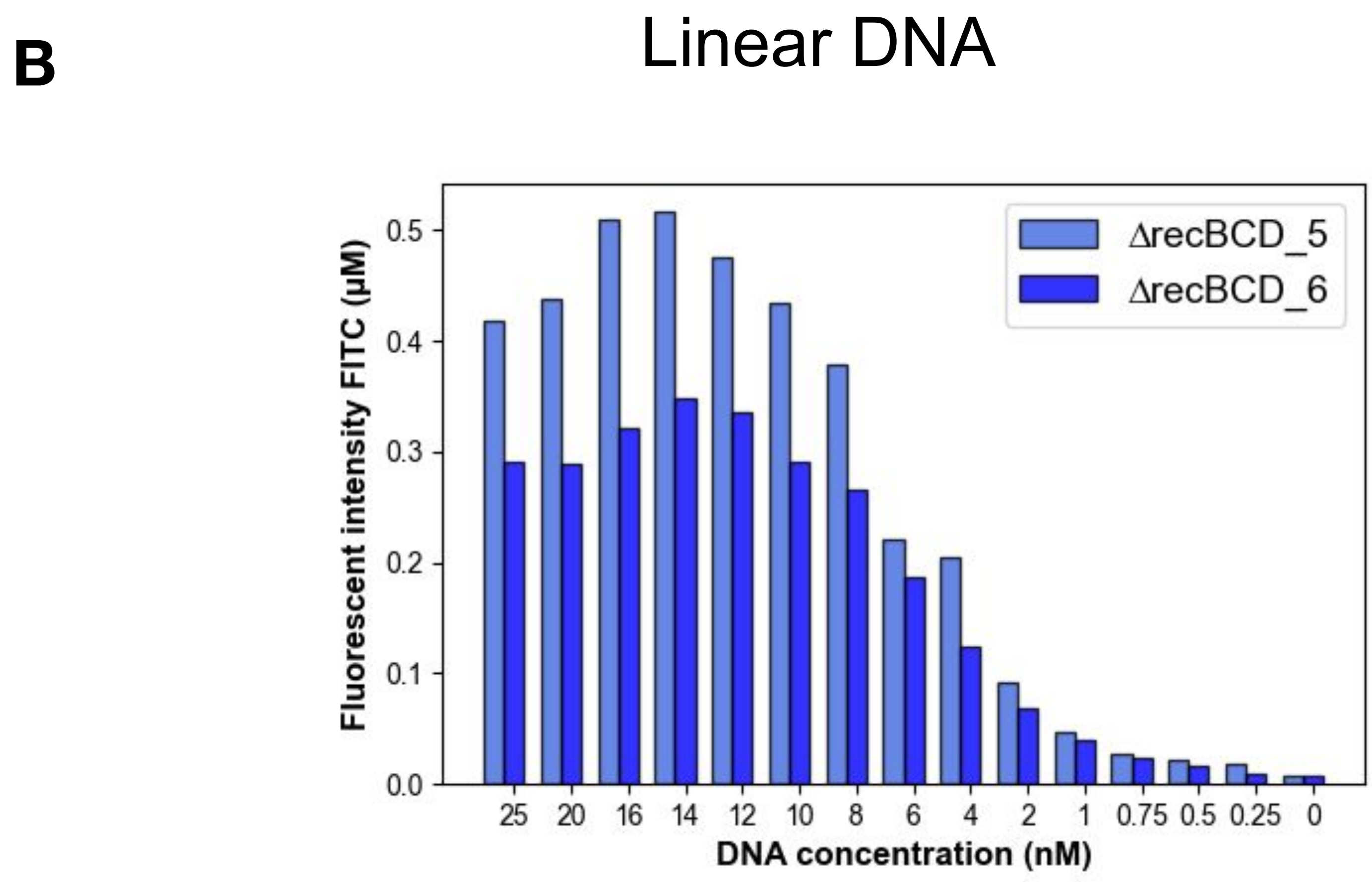


**Figure S4. Optimal linear and plasmid DNA concentration in Δ*recBCD* extracts**. (A) DNA calibration of plasmid DNA for two different batches of Δ*recBCD* CFS. (B) DNA calibration of linear DNA for two different batches of Δ*recBCD* CFS. To achieve high expression with less DNA, 10 nM was chosen for the rest of the experiments.

### Figure S5: Differential buffer optimization for *deGFP* at 30C (Δ*recB*/ Δ*recBCD* extracts from sonication)

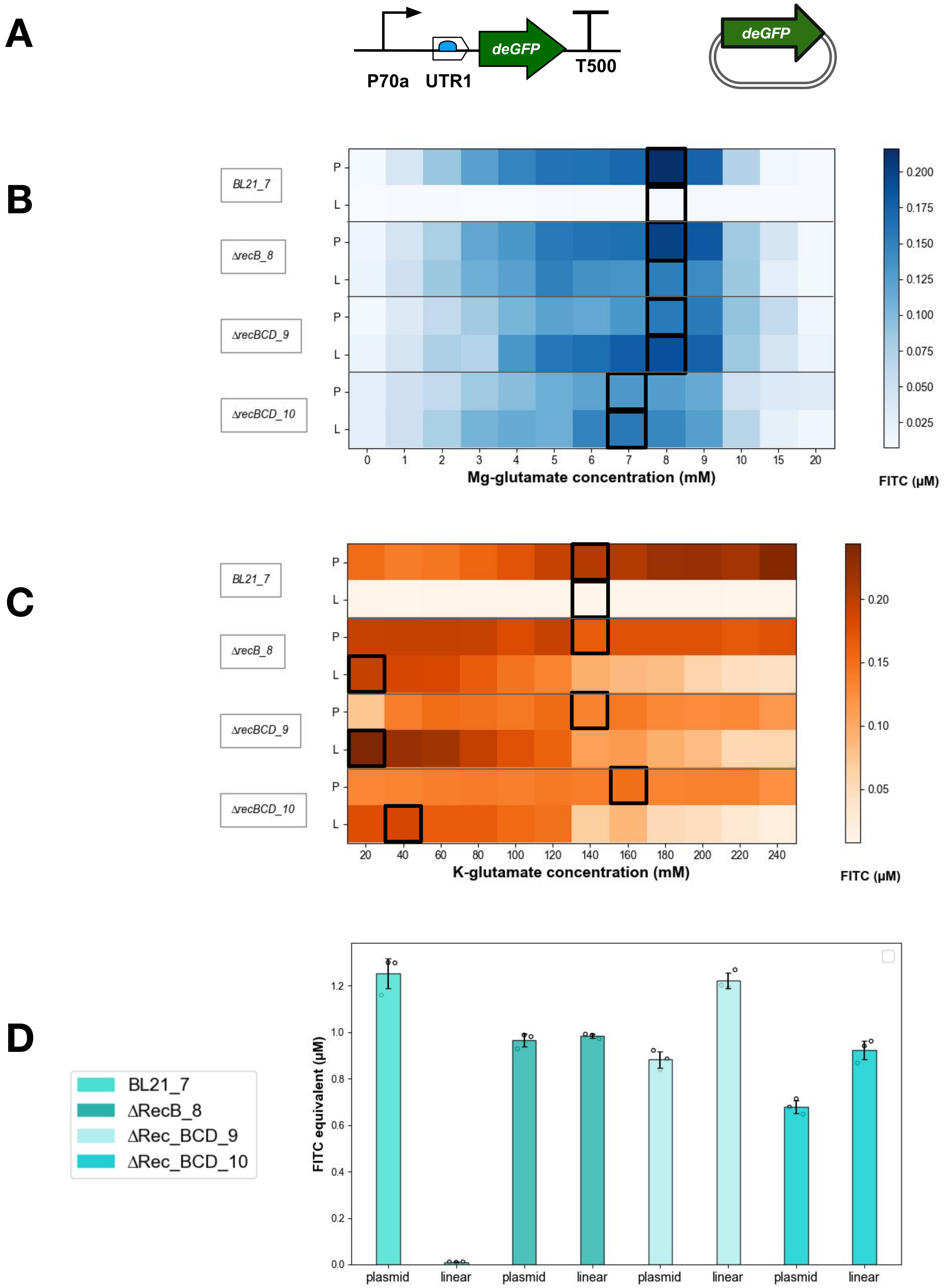


**Figure S5. Differential buffer optimization for *deGFP* at 30C (ΔrecB/ Δ*recBCD* extracts from sonication).** Buffer calibration for linear and plasmid DNA of *deGFP* reporter gene. (A) A 1361 bp linear DNA amplicon was amplified from plasmid p70a-deGFP and used for evaluating gene expression for buffer calibration. (B) Mg-glutamate buffer calibration was performed on lysates made by sonication with 1 nM of each DNA type using increasing concentrations from 0 to 20 mM (with K-glutamate fixed at 80mM). Square boxes indicate concentration points chosen. P= plasmid DNA, L= linear DNA. (C) After Mg-glutamate concentration was established for each lysate, K-glutamate was titrated from 20 to 270 mM. Squared boxes indicate concentration points chosen. P= plasmid DNA, L= linear DNA. Data shown in the heat-maps (B) and (C) are from a single experiment. (D) After buffer calibration, cell-free reactions were performed with 5 nM of each DNA type. Extract BL21 Rosetta2 (BL21_7) does not support linear DNA expression, whereas differentially optimized extracts from Δ*recB* (ΔrecB_8) and Δ*recBCD* strains (ΔrecBCD_9 and ΔrecBCD_10) exhibit linear DNA expression at near-plasmid levels. Data shown are mean±SD for 3 replicates done on the same day.

### Fig S6: Purification of linear DNA fragments after PCR is not required

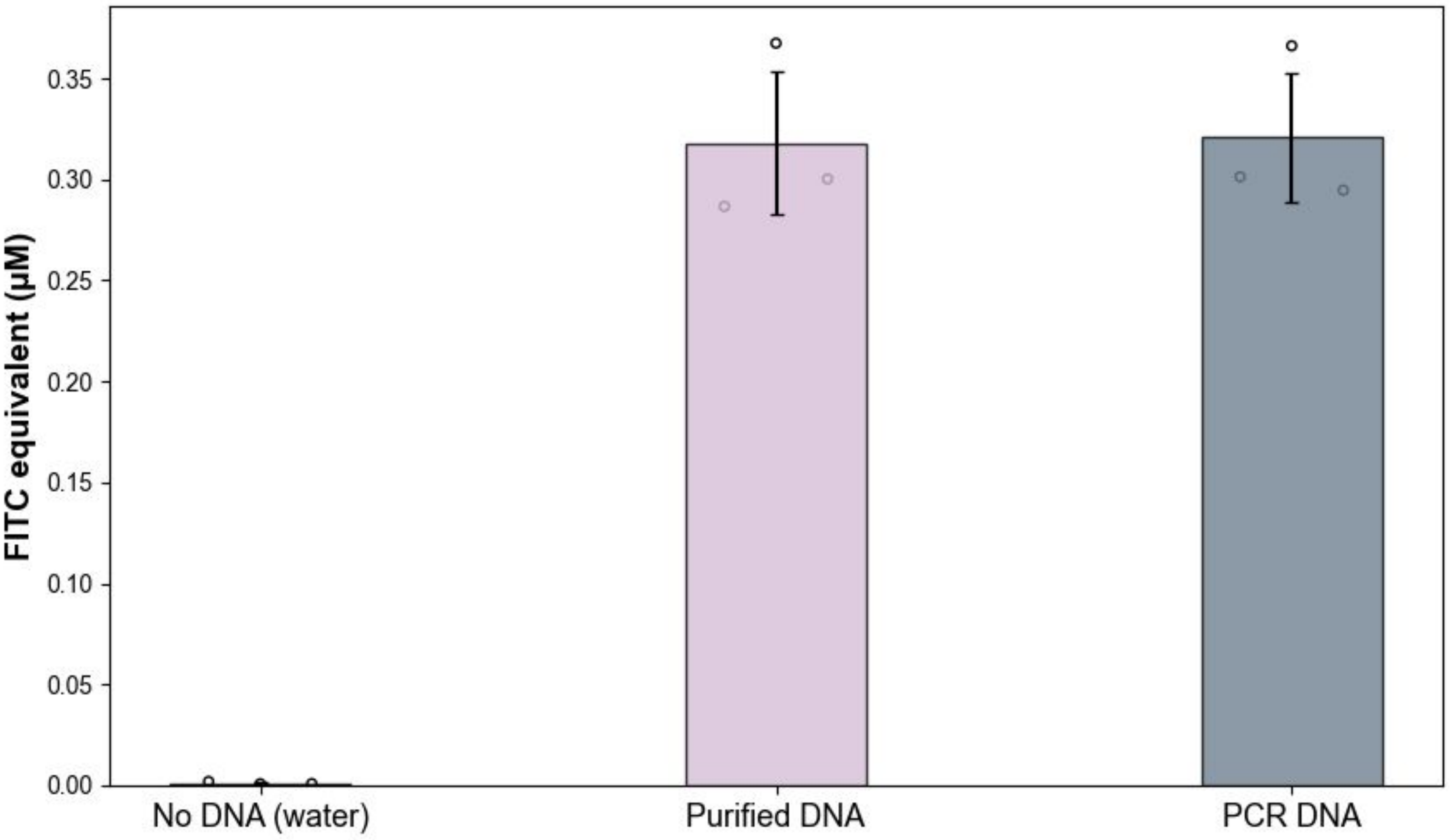


**Figure S6. Purification of linear DNA fragments after PCR is not required**. Equimolar amounts (10 nM) of linear fragments generated from PCR amplification with Invitrogen™ Platinum™ SuperFi™ polymerase were added directly to the cell-free mix, or after DNA purification. No significant difference in GFP expression was observed between them. Data shown are mean±SD for 3 replicates done on different days in ΔrecBCD_5 extract.

**Fig S7: Calibration of GamS and Chi inhibitors in BL21 CFS.**

A

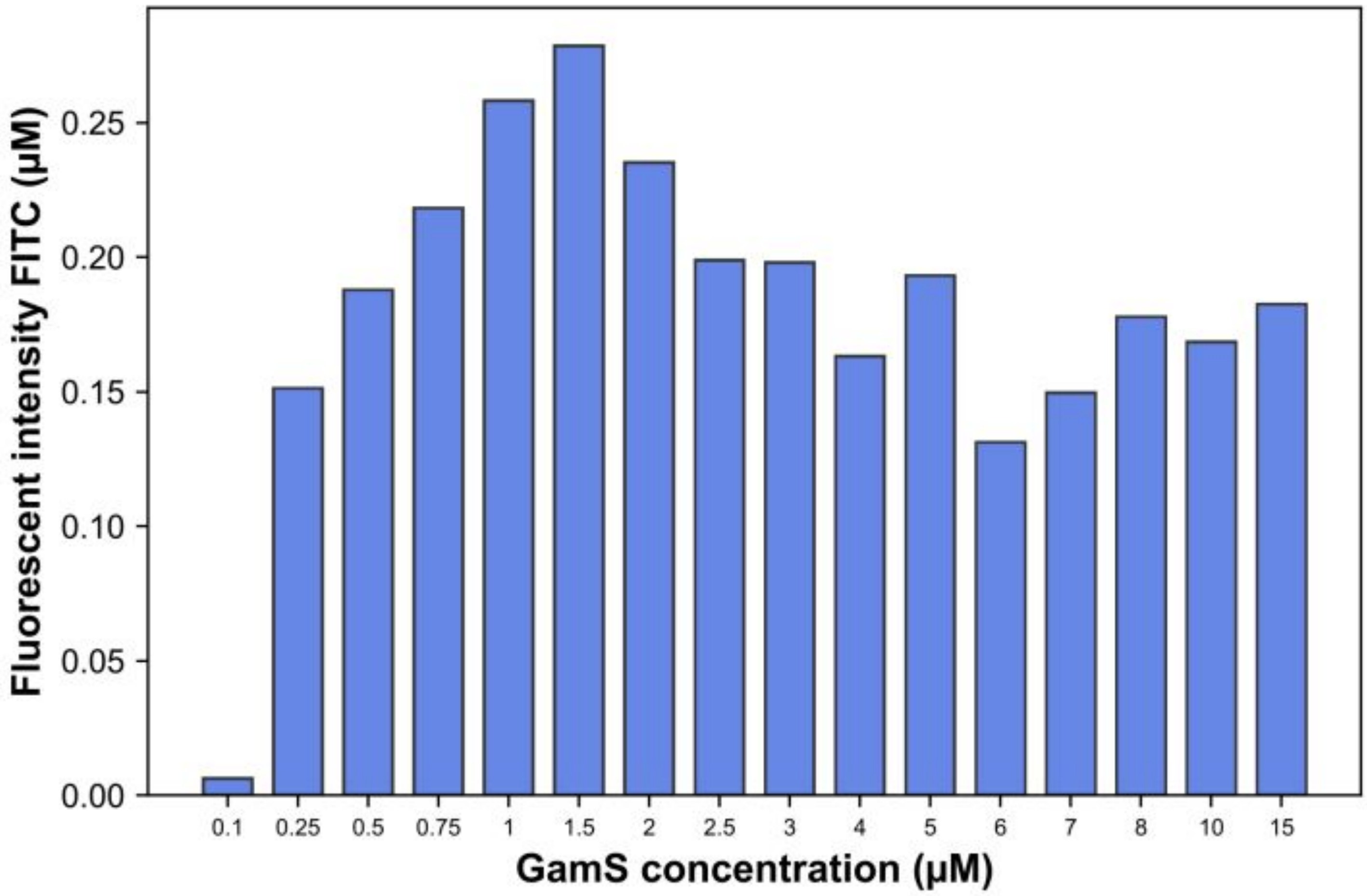


B

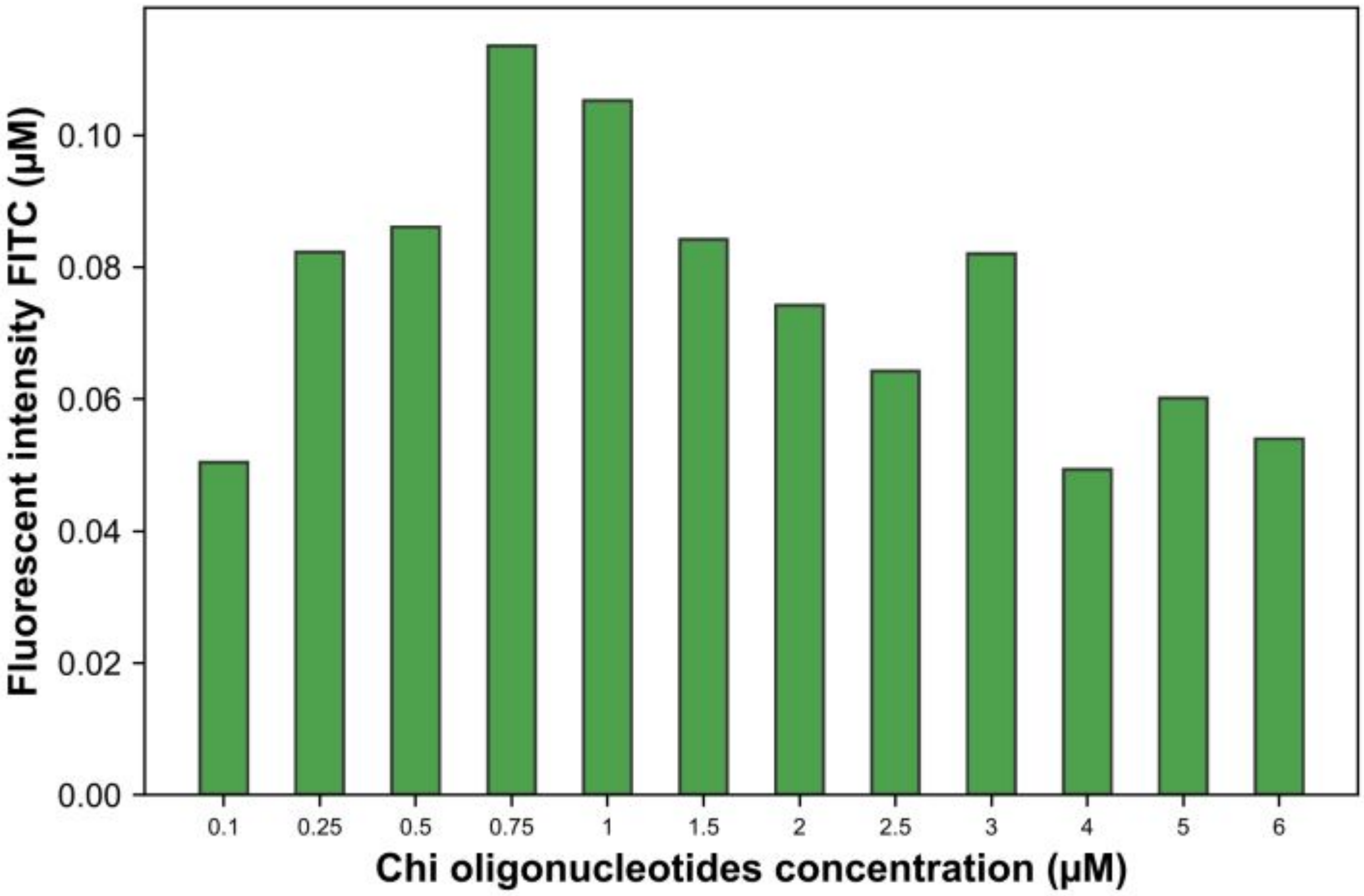


**Figure S7. Calibration of GamS and Chi inhibitors in BL21 CFS**. (A) GamS was produced and purified in-house (see Methods). The calibration was done at 10 nM of linear DNA. 2 μM of GamS was chosen as the working concentration. Data shown are from a single experiment in BL21_4 extract. (B) Chi6 oligos were annealed and added to the CFS according (see Methods). 1 μM of the Chi6 oligos was chosen as the working concentration. Data shown are from a single experiment in BL21_4 extract.

### Figure S8: GamS supplementation by doping/ pre-expression.

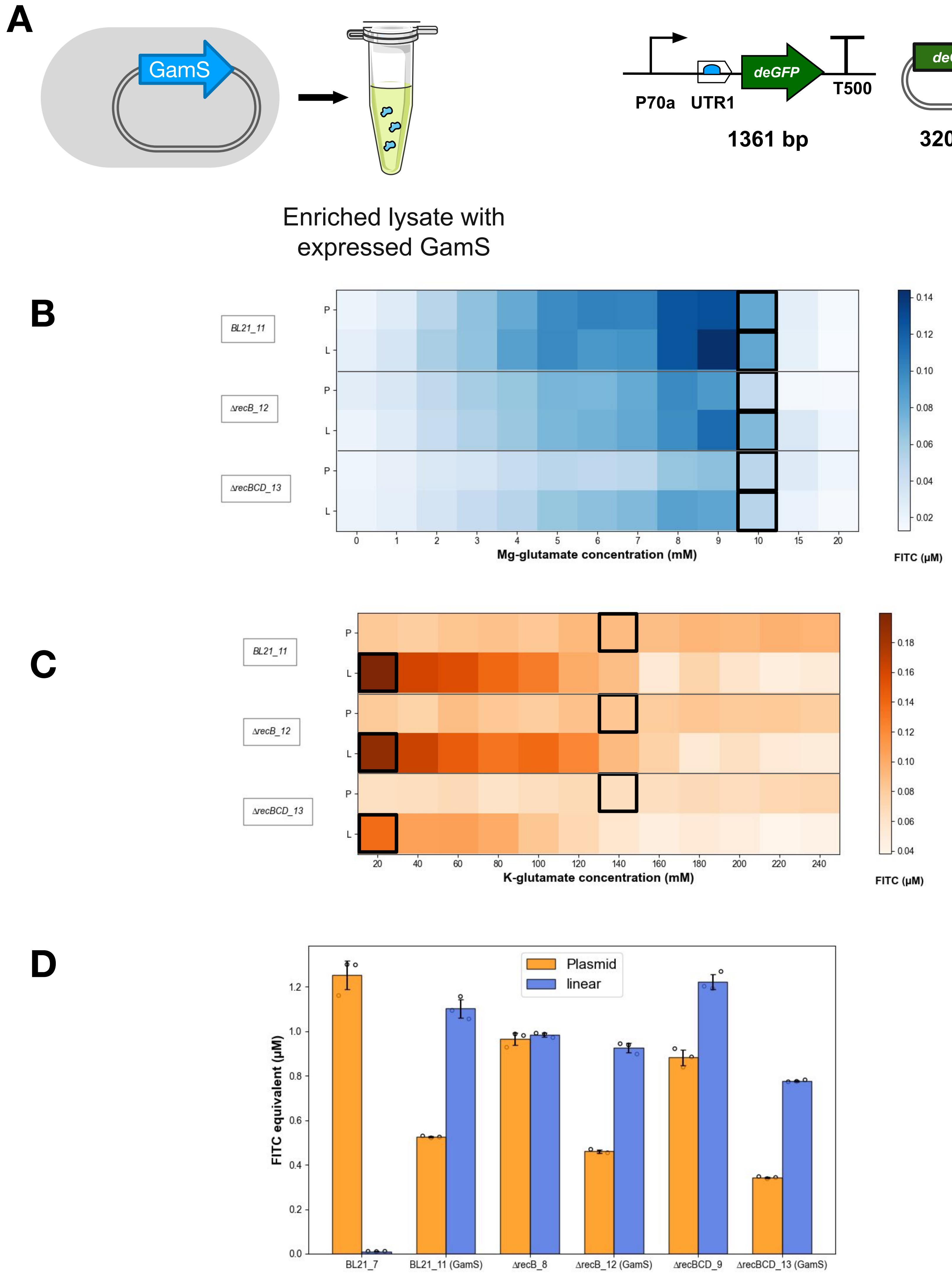


**Figure S8: GamS supplementation by doping/ pre-expression.** Buffer calibration for linear and plasmid DNA of *deGFP* in GamS-overexpressed lysates. (A) Parental BL21 Rosetta2 or modified (Δ*recB* and Δ*recBCD*) strains were transformed with GamS plasmid, and induced during expansion for lysate preparation. Linear or plasmid DNA, as used in Figure S5, were used as template for buffer characterization. (B) Mg-glutamate buffer calibration was performed on lysates made by sonication with 1 nM of each DNA type using increasing concentrations from 0 to 20 mM (with K-glutamate fixed at 80mM). Boxes indicate concentration points chosen. P= plasmid DNA, L= linear DNA. (C) After Mg-glutamate was established for each lysate, K-glutamate was titrated from 20 to 270 mM. Boxes indicate concentration points chosen. P= plasmid DNA, L= linear DNA. Data shown in the heat-maps (B) and (C) are from a single experiment. (D) After buffer calibration cell-free reactions were performed with 3 technical replicates with 5 nM of each DNA type. Lysate BL21 Rosetta2 GamS (BL21_11) allows expression from linear DNA in comparison to the wild-type BL21 Rosetta2 (BL21_7). Extracts from Δ*recB* strains GamS (ΔrecB_12) and/or Δ*recBCD* (ΔrecBCD_13) did not show an increase in linear DNA expression compared to extracts from their parental (ΔRecB_8 and ΔRecBCD_9) strains. The low K-glutamate concentration is also beneficial to linear DNA expression in these lysates. Data shown are mean±SD for 3 replicates done on the same day.

**Figure S9: Chemical modification of linear DNA ends in exonuclease containing cell-free extract.**

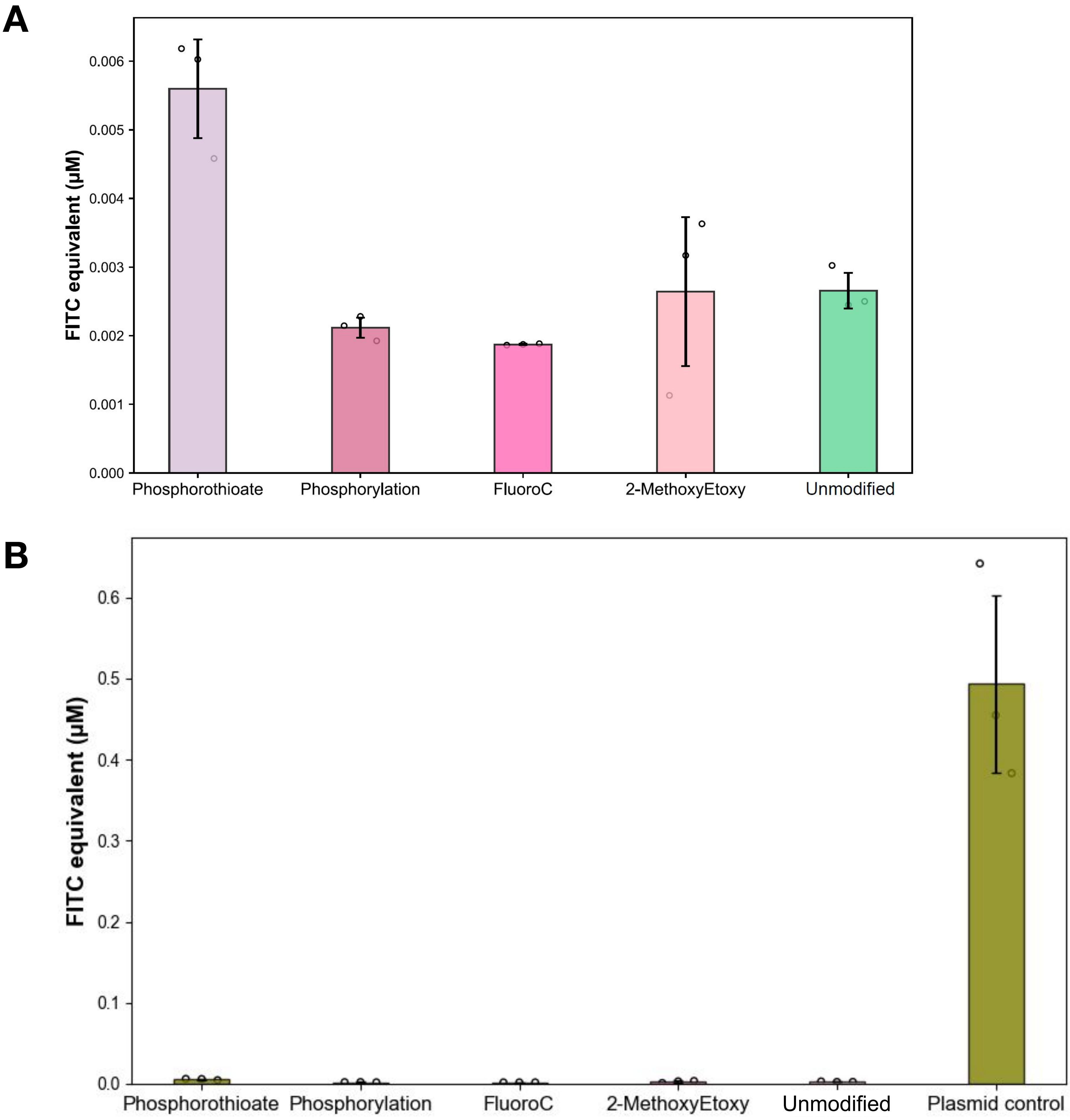


**Figure S9. Chemical modification of linear DNA ends in exonuclease containing cell-free extract. (A & B)** Assessment of the effect of various chemical modifications on the 3′ and 5′ ends of the linear DNA fragment on gene expression in BL21_4 extract. The reporter cassette was PCR amplified using various modified oligonucleotides and used as a template in cell-free reactions (Methods). Data shown are mean±SD for 3 replicates done on the same day. Additional experimental conditions are listed in Sup Table 6.

### Figure S10: Toehold-switch expression cassette.

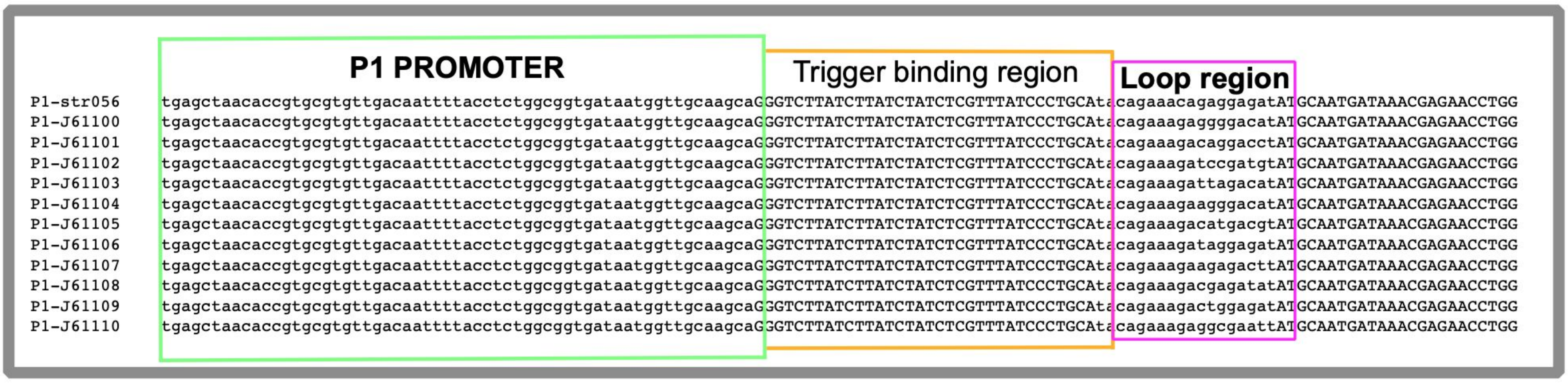


**Figure S10. Toehold-switch expression cassette.** Detail of aligned toehold variant sequences with the conserved P1 promoter sequence and trigger binding regions preceding the variable RBS containing hairpin loop region.

### Figure S11: HipO variant sequences

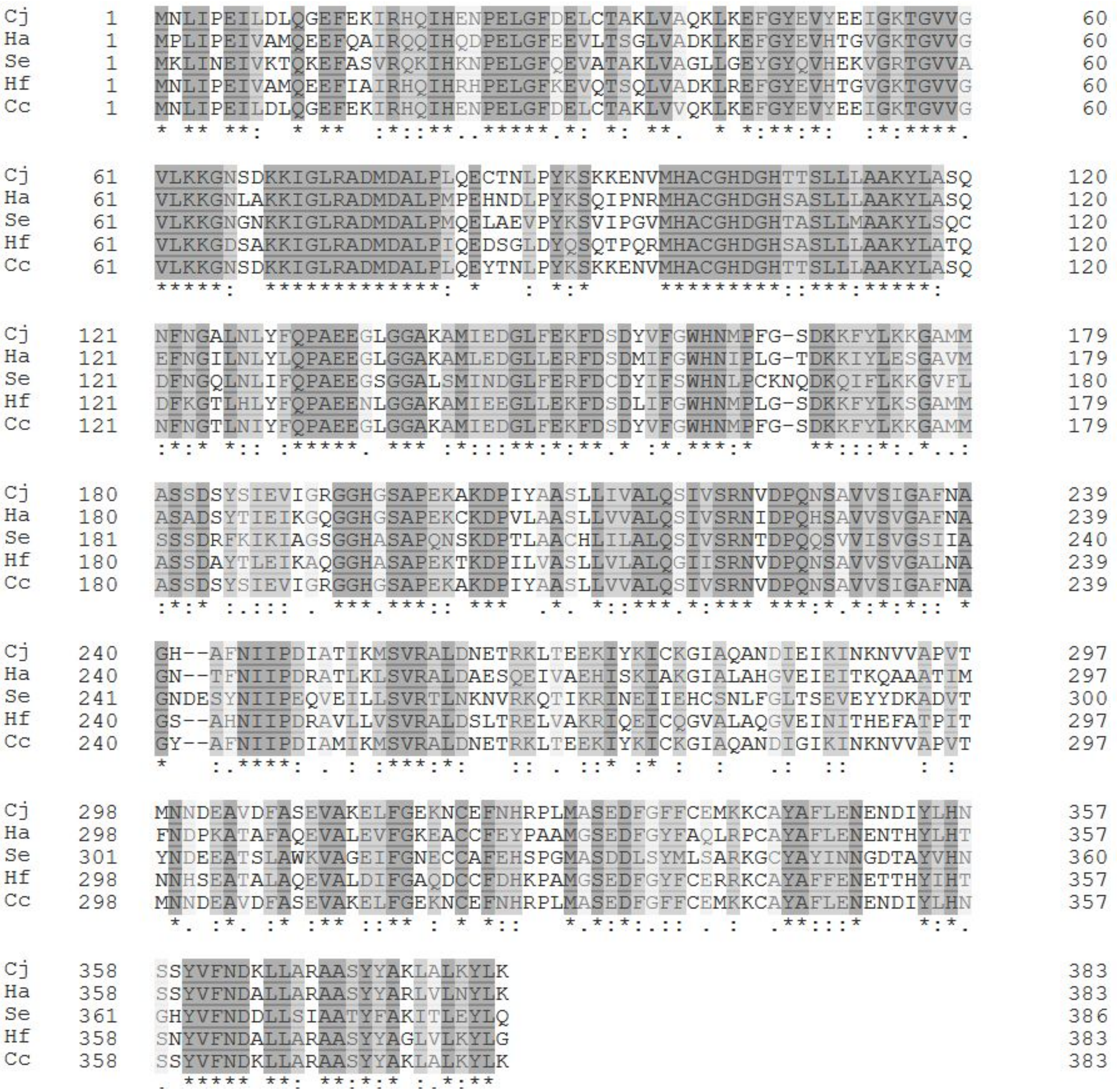


**Figure S11. HipO variant sequences**. Amino acid sequence alignment of HipO enzymes variants selected, synthesised and screened. The alignment was realised using Clustal Omega tool (https://www.ebi.ac.uk/Tools/msa/clustalo/). Identical amino acids are highlighted in dark grey, and similar amino acids in light grey.

#### Figure S12: Cell-free workflows time & cost calculation.

| | plasmid CF | | | GamS linear CF | | | ΔrecBCD linear CF | | |
|---|---|---|---|---|---|---|---|---|---|
| | STEP | COST | TIME | STEP | COST | TIME | STEP | COST | TIME |
| **Cell-free reaction** | Cell free reaction | 0.44€ / 20µl rxn | 10h | Cell free mix | 0.44€ / 20µl rxn | 10h | Cell free mix | 0.44€ / 20µl rxn | 10h |
| | | | | GamS | 0.72€ / 20µl rxn | | | | |
| **DNA preparation** | PCR amplification | -1.63€ Q5 PCR Mix<br>-10€ oligos<br>-1.8€ PCR cleanup mix | 3h | PCR amplification | -2.82€ Sfi PCR Mix<br>-10€ oligos | 2h | PCR amplification | -2.82€ Sfi PCR Mix<br>-10€ oligos | 2h |
| | Gibson cloning | 20€ kit | 16h | | | | | | |
| | colony check | -2 clones miniprep 4€<br>-2 clones sequencing | 2 days | | | | | | |
| | Maxiprep | 10€ | 1 day | | | | | | |
| **TOTAL** | | ≈ 51 € | 4-5 days | | ≈17,5 € | 12 h | | ≈14,5 € | 12 h |

**Figure S12. Cell-free workflows time & cost calculation.** Time and cost comparison of 3 methods for cell-free characterization of a genetic part: the traditional plasmid using cell-free, the GamS dependant linear DNA method and the developed method relying on the Δ*recBCD E. coli* strain. Costs are calculated for reagents, kits and outsourced services (oligo synthesis and plasmid sequencing) necessary to carry each workflow. Labor cost is not calculated but the duration of each process was estimated.
